# TIAR-dependent coordination of alternative splicing and lipid peroxidation is required for CML cell resistance to imatinib in the bone marrow stroma

**DOI:** 10.64898/2026.05.29.728710

**Authors:** Paulina Podszywalow-Bartnicka, Ewa Kozlowska, Remigiusz Serwa, Joanna Idaszek, Ewa Walejewska, Agnieszka Kepczynska, Anna Mietelska-Porowska, Paulina Pilanc-Kudlek, Magdalena Wolczyk, Leonard Schärfen, Bac Viet Le, Wojciech Swieszkowski, Katarzyna Piwocka, Tomasz Skorski, Karla M. Neugebauer

## Abstract

Chronic myeloid leukemia (CML) is treated with Abl1 tyrosine kinase inhibitors (TKIs). Quiescent cancer cells residing in the bone marrow (BM) can survive the treatment and cause CML relapse. We previously found that a subset of alternative splicing (AS) changes detected in CML cells surviving months of therapy are initiated within hours of treatment onset. Here, we investigated how AS in CML cells is modulated by the human BM microenvironment. By incorporating humanized BM niche models *in vivo,* we uncovered stroma-induced transcriptome adaptation that influences transcriptional regulation, transmembrane transport, lipid metabolism, the tricarboxylic acid cycle, and respiratory electron transport. We identified RNA-binding protein TIAR (T-cell intracellular antigen-related protein) as a key mediator of CML survival under TKI imatinib treatment. Our data show TIAR-dependent coordination of RNA processing with the metabolic program induced by stromal interaction. Quantitative nascent proteome analysis revealed that TIAR silencing affects the synthesis of metabolic enzymes and proteins involved in imatinib-induced erythroid differentiation. Besides, TIAR knockdown increased lipid peroxidation in untreated cells and decreased reduction potential in cells upon imatinib treatment. Taken together, TIAR deficiency reduces CML survival, possibly by inducing ferroptosis. These findings identify TIAR-dependent RNA processing within the BM niche as a previously unrecognized mechanism of CML therapy resistance and a potential therapeutic vulnerability.

## INTRODUCTION

The extracellular (micro) environment profoundly shapes the activity of intracellular processes that govern cell survival. Nutrient availability, ionic balance, and the presence of signaling molecules collectively determine critical cellular parameters, including energy state and gene expression. Thereby, the microenvironment of cancer cells defines the cellular response to therapy. Drugs effective against actively dividing cells in the blood circulation frequently lose efficacy against dormant, microenvironment-sheltered cells. This poses a challenge in the treatment of cancers harboring hypoxic regions. Cell quiescence is a key pro-survival state in acute and chronic myeloid leukemias (AML and CML) that originate from hematopoietic myeloid progenitor cells and develop within the bone marrow (BM). CML cells acquire dormancy in response to hypoxia (*1–3*) and to interaction with BM stromal cells (*4, 5*) (for review see (*6*)). Premature discontinuation of therapy results in CML recurrence in up to 98% of cases, even in patients who have achieved molecular remission (*7*). The propagation of TKI-resistant cells drives progression to blast crisis, the most aggressive stage of the cancer. Therefore, it remains a critical unmet challenge to determine the environment-activated mechanisms that sustain cancer cell survival (*2, 8*).

Various experimental setups have been developed to investigate how components of the hypoxic BM niche influence myeloid leukemia cell biology. In *ex vivo* 2D and 3D co-culture models pairing leukemia cells with BM stromal cells, such as mesenchymal cells or fibroblasts, influences DNA damage repair (*9*), mRNA translation (*10*), and cellular metabolism (*11, 12*). BM stromal fibroblasts modulate leukemic cell behavior through both secreted factors (*9, 13–15*) and direct cell contact-dependent mechanisms, including adhesion (*16, 17*), tunneling nanotubes (TNTs) (*11*), and Delta-Notch signaling (*4, 18, 19*). Despite the limited cellular diversity inherent to this system, multiple studies have consistently demonstrated that CML cells acquire TKI insensitivity in this *ex vivo* setup (*13, 14, 16, 17, 20–23*), driven by non-genomic cellular changes. *In vivo,* xenotransplantation of CML cells together with human BM mesenchymal stem cells (hMSC) into immunocompetent mice conferred protection against TKI treatment, a response shown to be stimulated by interleukin 17 (*24*).

The cellular environment dictates the profile of proteins synthesized within cells. This is coordinated by molecular mechanisms regulating gene expression at the steps of transcription activation and RNA processing. The synthesis of new protein variants through alternative splicing in CML has emerged as a mechanism supporting malignant progression (*25*), providing protection from apoptosis (*26*), and mediating early CML cell response to TKI treatment (*27*). The importance of RNA-binding proteins (RBPs), which orchestrate RNA processing at multiple levels, is therefore particularly compelling in the context of microenvironment regulation. We previously demonstrated that RBPs selectively coordinate protein synthesis in CML cells, in an extracellular condition-specific manner (*10*). We found that TIAR, encoded by TIA1 cytotoxic granule associated RNA binding protein like 1 (*TIAL1*) gene, is required for leukemia cell survival following translation inhibitor treatment under hypoxic conditions but not under normoxia (*10*). In response to cellular stress, TIAR is recruited to cytoplasmic stress granules and nuclear speckles involved in co-transcriptional RNA processing (*28, 29*). As one of the AU-rich element-binding RBPs, TIAR modulates alternative splicing (*30–33*) and translation accessibility of target transcripts (*10, 34–37*). TIAR activity influences cell cycle progression and promotes cell quiescence (*32, 38, 39*). Collectively, TIAR is an important component of the downstream signaling of the integrated stress response, a pathway that promotes TKI resistance in CML (*40, 41*). However, most of the current knowledge of TIAR function comes from experiments performed under atmospheric oxygen conditions, leaving the significance of TIAR activity, including its impact on alternative splicing, largely unknown in the context that is physiologically relevant for human leukemia therapy resistance.

Here, we demonstrate that TIAR activity is critical for leukemia cell survival in the BM under treatment with imatinib (IM), a TKI used as a first-line drug in CML. Our data position TIAR within a regulatory hub that coordinates RNA processing and lipid peroxidation with microenvironment conditions. Additionally, we present several *in vivo* experimental systems designed to interrogate treatment response and transcriptome adaptation of CML cells within the human BM stromal microenvironment.

## RESULTS

### Human CML cells survive imatinib treatment in the bone marrow stroma

To model the interaction of CML cells with BM stroma, we adopted an experimental framework originally developed for AML research (*42*). Ossicle-like xenotransplants (hereafter referred to as ‘ossicles’) were generated by subcutaneous transplantation of human BM mesenchymal stem cells from healthy donors (hMSC) into immunocompromised mice. As the principal progenitors of the major BM stromal components, including osteoblasts, adipocytes, and chondrocytes, hMSCs provide a physiologically relevant stromal niche (*43*) (Fig. 1A). To promote ossification, mice were administered human parathyroid hormone (PTH) (*42*). Then, Lin-CD34+ cells isolated from CML patient samples were injected and colonized the ossicles. Following 12 weeks of engraftment, mice underwent a 2-week course of IM treatment, consistent with our previously established protocol in humanized mouse models (*44*).

**Figure 1.**
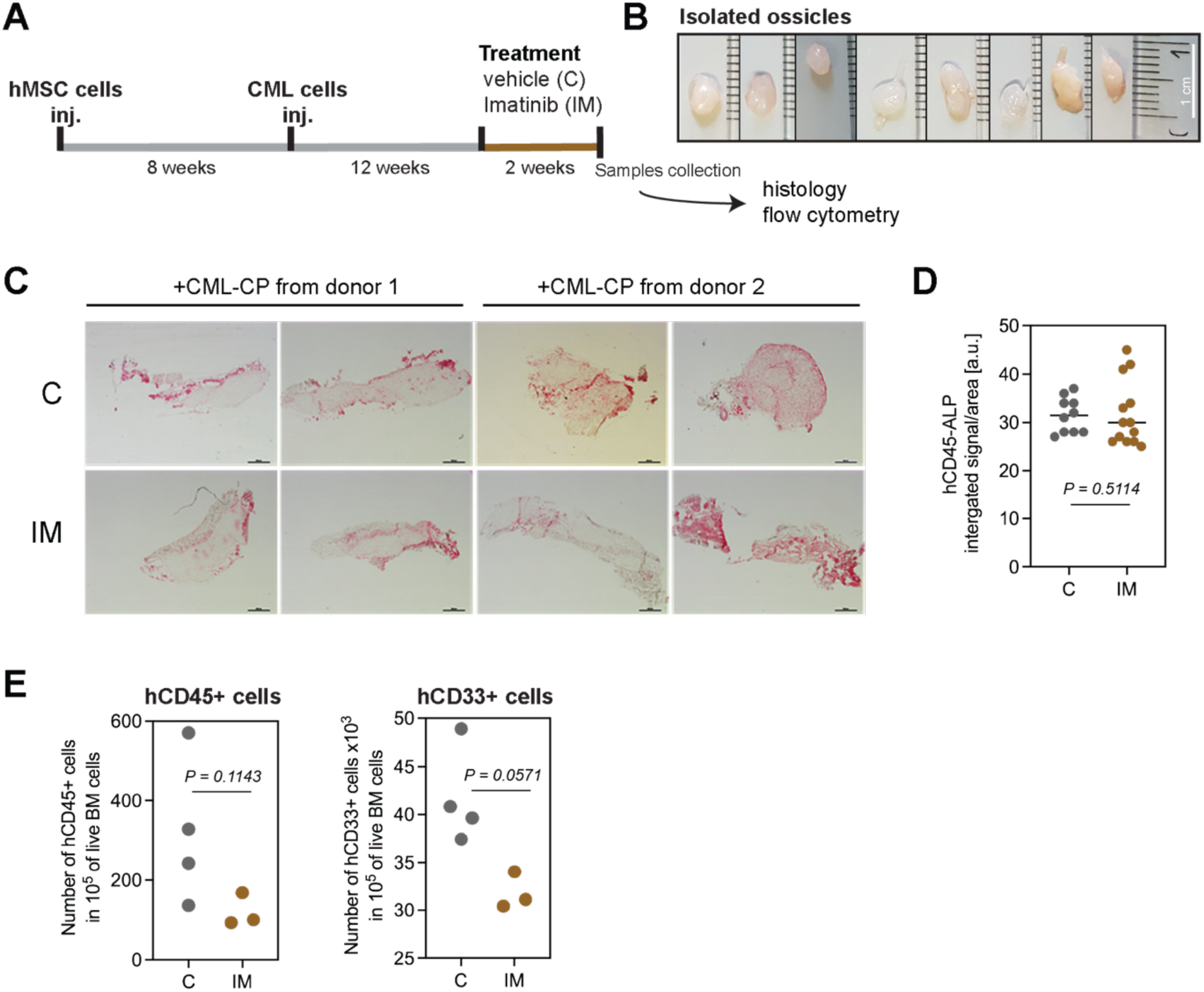
Leukemia cells survive imatinib treatment in ossicle-like xenografts. **(A)** Schematic explanation of the experiment. Following the period of 8 weeks of ossicle formation by hMSCs, the human chronic myeloid leukemia (CML-CP) primary Lin-CD34+ cells were injected, and after another 8 weeks, treatment with imatinib (IM) or vehicle (C) was initiated for 2 weeks. At the end, the ossicles were isolated for histological analysis, and mouse tissue samples were collected. **(B)** Representative ossicles photographed with a Samsung mobile phone camera along a ruler (cm) for size reference; white scale bar = 1cm. **(C)** Images of ossicle specimens after immunohistology detection of leukemia cells (from two different donors) with antibody against human CD45 and following alkaline phosphatase (ALP) reaction (visible in red) taken under Nikon Ni upright microscope with Nikon Plan UW 2x/0,06 objective in brightfield; scale bar = 1 mm. **(D)** Quantification in Fiji of ALP signal within the tile-scanned images of ossicles in **(C)**. Signal intensity normalized to the area of an ossicle in each specimen analyzed is represented by a dot; 2-3 consecutive sections through the middle of each ossicle from C and IM-treated mice (n=5 ossicles in each treatment) were analyzed. **(E)** Number of cells detected with fluorochrome-conjugated antibody against human CD45 (hCD45+) or CD33 (hCD33+) in the sample of mouse bone marrow (BM) from the tibias (n=4 and 3 in C and IM, respectively). Results of flow cytometry analysis were normalized to the number of live cell events recorded in each sample (provided on the y-axis). **(D-E)** The two-tailed, nonparametric Mann-Whitney test was used to compare results from C vs IM, and the exact significance (*P*) values are presented in the graphs.

The isolated ossicles measured approximately 0.5 mm in diameter and presented a uniform morphology (Fig. 1B). Histological analysis revealed no bone-like tissue organization. Instead, ossicles contained clusters of leukemia cells exhibiting increased hematoxylin staining attributable to a high nucleus to cytoplasm ratio. Immunostaining confirmed that these clusters were negative for the mouse and human CD31 protein, a marker of endothelial cells, and positive for the pan-human hematopoietic marker CD45 (hCD45) (Fig. 1C, S1). Quantification of hCD45-positive cells in cryosections revealed no significant difference in CML cell abundance between ossicles from vehicle-and IM-treated mice (Fig. 1D). Simultaneously, IM treatment reduced the numbers of hCD45-and hCD33-positive (human myeloid-biased) cells in the BM from mouse tibiae by approximately 62% (199.2 ± 111.4) and 28% (9.8 ± 3.1), respectively (Fig. 1E). Altogether, these findings demonstrate that CML cells can successfully home to human ossicles and persist in organized cell clusters that are refractory to IM treatment.

### Human bone marrow mesenchymal cells induce profound transcriptional reprogramming of CML cells in a xenograft model

To investigate the impact of human bone marrow stromal cells on leukemia cell transcriptome with greater experimental control over cell numbers, we employed a subcutaneous xenograft model. To this end, leukemia cells suspended in Matrigel were implanted either alone (mono) or co-implanted with primary hMSCs (Fig. 2A). We utilized the K562 cell line, which is derived from a CML patient in blast crisis and widely used as a CML model. The cells were lentivirally transduced with a construct encoding green fluorescent protein (GFP) and luciferase (K562/luc), and a non-targeting shRNA, which served as the reference in subsequent experiments (shNEG). Xenografts were growing for two weeks. Co-implantation with hMSCs increased mean xenograft weight by about 1.5-fold compared to mono, suggesting that K562/luc cells were the dominant contributor to the xenograft mass (Fig. 2B). Stronger Sirius Red staining of tissue sections revealed enhanced collagen deposition in hMSC-containing xenografts compared to mono (Fig. S2A), and a similar proportion of GFP-positive cells between conditions (Fig. S2B).

**Figure 2.**
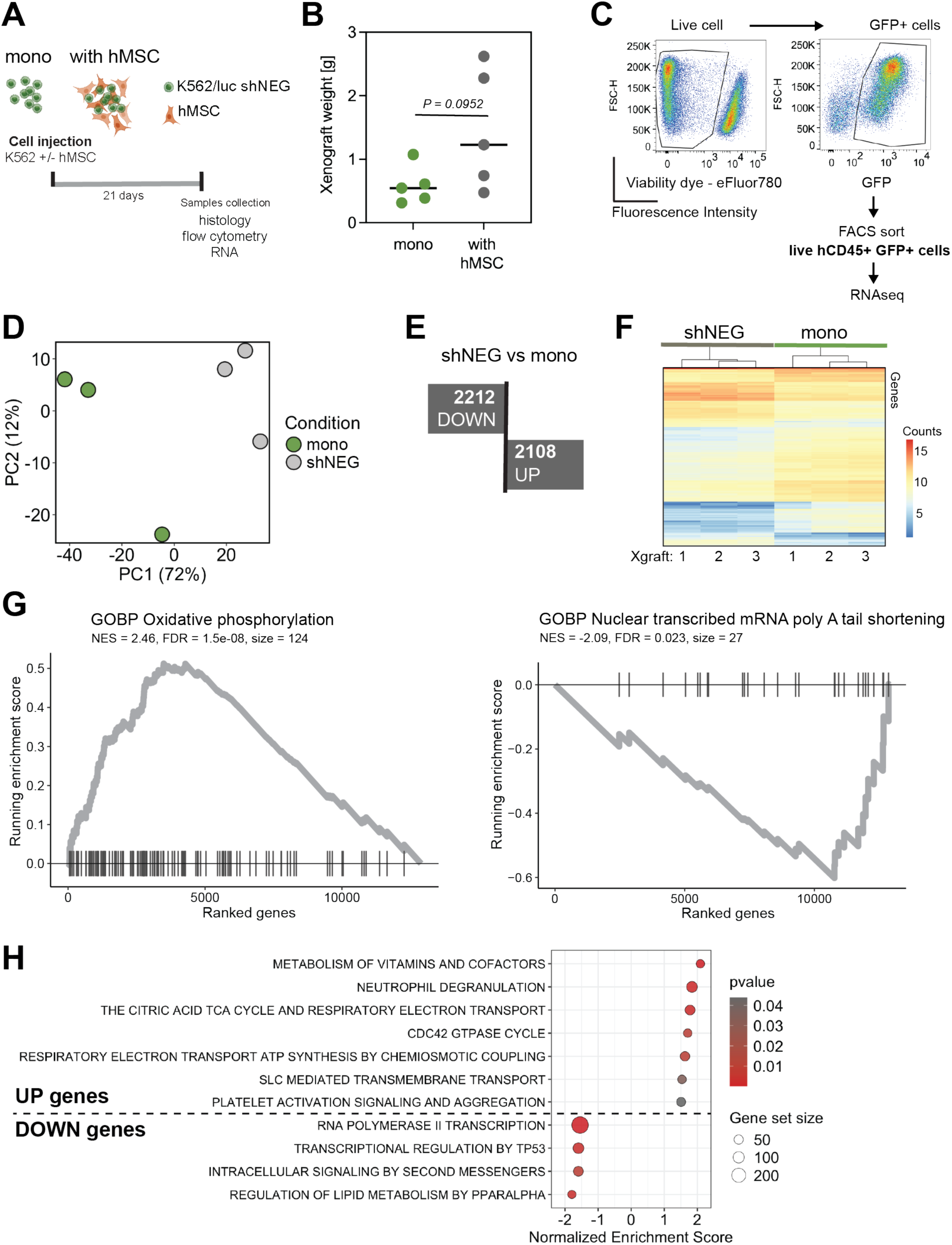
**Interaction with human bone marrow stromal cells changes the transcriptome of leukemia cells xenotransplanted *in vivo*** Results from K562/luc (expressing shNEG, Firefly luciferase, and GFP) alone (mono) or mixed with human primary bone marrow mesenchymal stem cells (with hMSC) were subcutaneously injected in mice to form xenografts. **(A)** Scheme demonstrating the experimental setup; samples were collected 21 days after subcutaneous injection of cells in mice. **(B)** Weight of xenografts formed by K562/luc mono or a mix with hMSCs in mice (n=5 in each variant), and the two-tailed, nonparametric Mann-Whitney test was used to check the difference significance; the exact significance (*P*) values are presented in the plot. **(C-H)** Short-read sequencing RNA analysis results of samples from cells isolated from xenografts mono and with hMSC treated for the last 14 days with vehicle (shNEG) by the fluorescence-activated cell sorting (FACS) with BD Aria. **(C)** Representative scatterplots (n=3) with the gating strategy used for the FACS-sorting of live xenograft cells (negative for Viability dye-eFluor780 staining), expressing GFP (GFP+) and stained positive with the antibody against human CD45 with BV421 fluorophore (hCD45+). **(D-G)** Results of differential expression analysis of the variant shNEG versus mono by DeSeq2 of RNA sequenced from FACS-sorted cells. Presented are the results from cells isolated from three different xenografts for each variant. **(D)** Principal component analysis. **(E)** Number of genes with fold change absolute value ≥ 50% and p-value corrected for multiple testing using the Benjamini-Hochberg P_adj._ ≤ 0.05, that are upregulated (UP) or downregulated (DOWN) in cells from xenograft (Xgraft) shNEG versus mono. **(F)** Expression level (counts corrected for the sample sequencing depth) in each xenograft (n=3) of the top 150 genes (each row) with the most significant (P_adj._) fold change in expression level.**(G)** The top of Gene Ontology Biological Processes (GOBP) terms from the Gene Set Enrichment Analysis (GSEA) of the DeSeq2 results, with the highest positive (left) and negative (right) normalized enrichment score (NES), *P_adj._* ≤ 0.05 (FDR) and number of genes in the sample annotated per term (size) > 20. The running enrichment score is presented by a grey line for each of the ranked genes marked with a vertical black line. **(H)** The Reactome terms identified by the GSEA of the UP or DOWN genes in the variant shNEG versus mono, with NES ≥ 1.5 (absolute value), p value ≤ 0.05, and size > 30.

This experimental setup enabled the isolation of live human leukemia cells co-expressing GFP and hCD45 from xenografts by flow cytometry-activated cell sorting (FACS)(Fig. 2C, S2C). The absolute number of leukemia cells in the hMSC-containing xenografts in vehicle-treated animals was comparable to that from the mono variant (Fig. S2B, S2C). To assess stromal influence on leukemia cell gene expression, we compared the transcriptome of K562/luc cells isolated from shNEG to those from mono xenografts. Short-read sequencing of polyadenylated transcripts (mRNA) revealed extensive transcriptional differences as evidenced by principal component analysis of normalized gene expression data (Fig. 2D), with 2,108 genes significantly upregulated and 2,212 downregulated in shNEG relative to mono xenografts (Fig. 2E). These changes were reproducible across individual xenografts within each condition (Fig. 2F).

Functional enrichment analysis of differentially expressed genes identified significant upregulation of proteins involved in oxidative phosphorylation (Fig. 2G), translation regulation (Fig. S3A), and SLC-mediated transmembrane transport, as well as downregulation of proteins mediating mRNA poly(A) tail deadenylation (Fig. 2G). Pathway-level analysis of upregulated genes revealed enrichment in metabolic processes (vitamins and cofactors metabolism, the tricarboxylic acid (TCA) cycle, mitochondrial respiratory electron transport), alongside pathways governing vesicle-mediated signaling (neutrophil degranulation, platelet activation), and actin cytoskeleton polymerization (CDC42 GTPase cycle) (Fig. 2H, S3B). Conversely, downregulated genes were enriched in processes related to transcriptional regulation (RNA Polymerase II-mediated transcription, TP53-dependent transcriptional control), intracellular signal transduction, and lipid metabolism (Fig. 2H, S3B). Taken together, these findings demonstrate that the presence of human BM stromal cells drives broad metabolic and transcriptional reprogramming in CML cells, consistent with previously reported observations in co-culture systems (*12*).

### Imatinib-induced transcriptome changes in the xenograft model recapitulate gene expression patterns observed in CML patient bone marrow biopsies

To assess the transcriptional impact of IM, mice bearing hMSC-containing K562/luc shNEG xenografts were treated for 2 weeks (Fig. 3A). IM treatment reversed a substantial portion of transcriptomic changes induced by hMSC co-implantation (Fig. S4). To contextualize these findings clinically, we compared transcriptional response to IM in xenograft-derived leukemia cells (shNEG) to that of CD34-positive leukemia cells isolated from BM biopsies of a CML patient obtained at diagnosis and following 6 months of IM therapy (Fig. 3B). This analysis identified a set of genes consistently upregulated (n=762) and downregulated (n=755) upon IM treatment in both datasets (shNEG & CML) (Fig. 3B), revealing a pattern of reversed regulation of processes previously shown to be modulated by hMSC co-implantation (Fig. 2G, 2H, S3, S4). These processes were related to transcription regulation, transmembrane transport, lipid metabolism, TCA, and respiratory electron transport (Fig. 3C). To interpret relevance of such changes in a broader perspective we examined the expression of leading-edge genes in a published (*45*) single-cell RNA sequencing dataset derived from BM biopsies of healthy donors (8 patients) and CML patients, categorized by clinical response to IM therapy: group A (9 patients) - responded to IM within 12 months; group B (9 patients) – IM treatment failed within 18 months but responded to other TKIs; group C (6 patients) – resistant to IM and other TKIs (Fig. 3D). Most of the genes upregulated following IM treatment in the shNEG & CML comparison were also progressively upregulated in CML patients with increasing IM resistance relative to healthy donors and IM-responsive patients (Fig. 3D). The corresponding trend among downregulated genes was less consistent. Together, these findings suggest that the hMSC-containing xenograft model partially recapitulates the transcriptional context and stromal influence experienced by CML cells within the human BM microenvironment.

**Figure 3.**
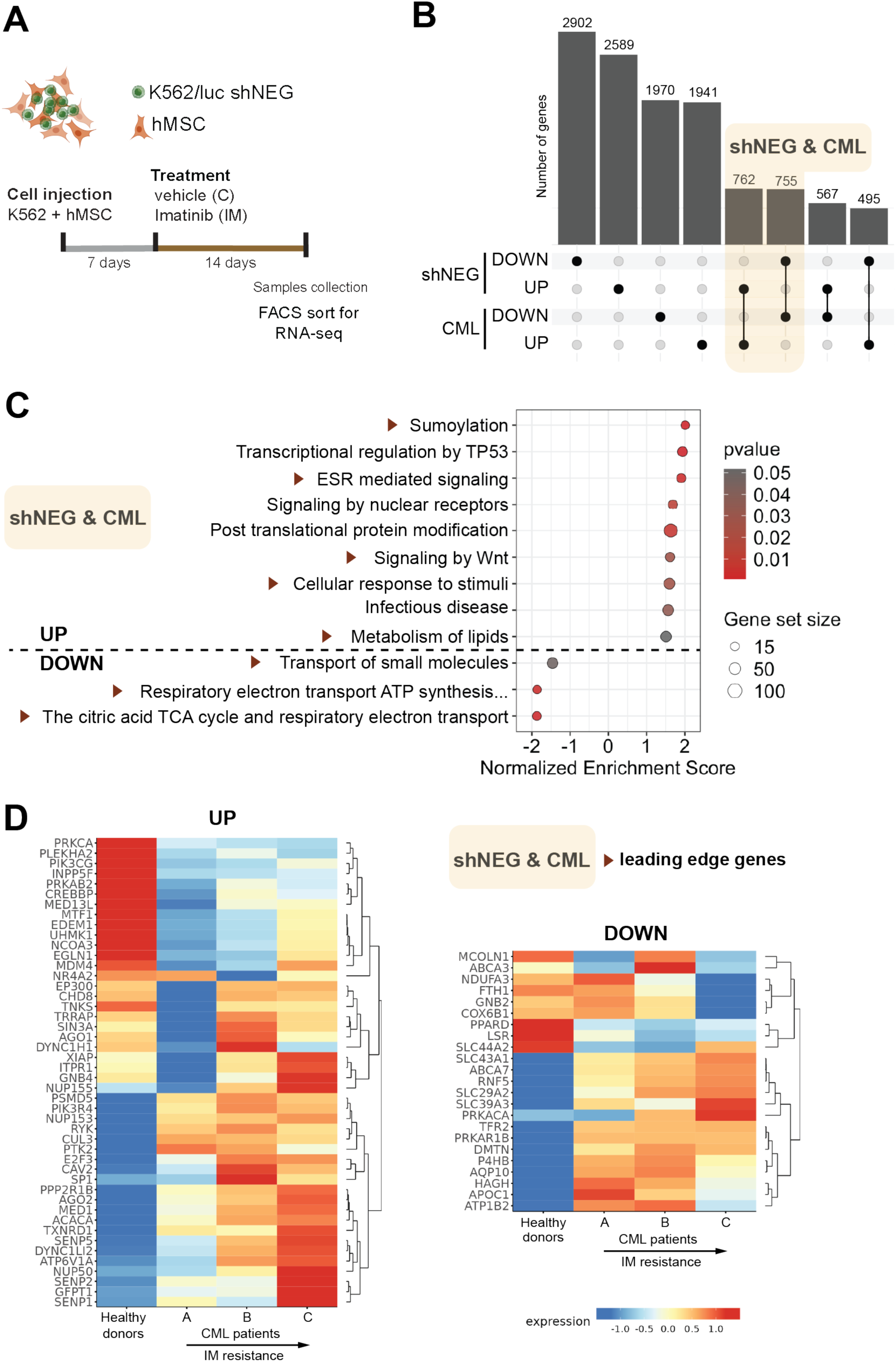
Imatinib treatment alters the transcriptome of CML cells residing in human bone marrow stroma. **(A)** Scheme explaining experimental steps of xenograft formation by K562/luc (expressing shNEG, Firefly luciferase, and GFP) mixed with human primary bone marrow mesenchymal stem cells (hMSC) subcutaneously injected in mice 7 days before initiation of treatment with IM for the following 14 days, when the GFP and human CD45 positive cells were FACS-sorted from xenografts for RNA isolation and sequencing. **(B)** Number of intersecting genes (compared groups indicated by black dot) that at the RNA level are upregulated (UP; log2 value of fold change ≥ 0.6) or downregulated (DOWN; log2 value of fold change ≤-0.6) in shNEG cells xenografts imatinib versus vehicle treated and in CD34 positive cells from bone marrow biopsies of a patient with CML (data deposited at GEO under accession number GSE310243; from (*27*)) obtained after 6 months of imatinib therapy (combined samples SRR36072325 and SRR36072321) versus obtained at the diagnosis (combined SRR36072320 and SRR36072324). **(B-D)** Yellow shadow marks a group of genes that are UP or DOWN upon imatinib treatment in shNEG and CML cells (shNEG&CML), selected for subsequent analysis in **(C)** and **(D). (C)** Reactome terms identified by the GSEA analysis of the UP or DOWN genes in shNEG&CML with normalized enrichment score ≥ 1.5 (absolute value), p value ≤ 0.05, and number of genes in the sample annotated per term (size) ≥ 15. **(C-D)** The Reactome leading-edge genes identified by GSEA, annotated to terms marked with a brown triangle in **(C)** were selected to compare their expression level in single-cell RNA-seq data in **(D)** from bone marrow biopsies of healthy donors and CML patients with different responses to imatinib therapy. **(D)** Analysis of data at the Single-cell atlas of diagnostic Chronic Myeloid Leukemia bone marrow (scdbm) for CD34+ cells subtype (*45*). Data and detailed description of patient classification available: http://scdbm.ddnetbio.com; A – responded to IM within 12 months; B – IM treatment failed within 18 months; C – resistant to IM and other TKI-s.

**Figure 4.**
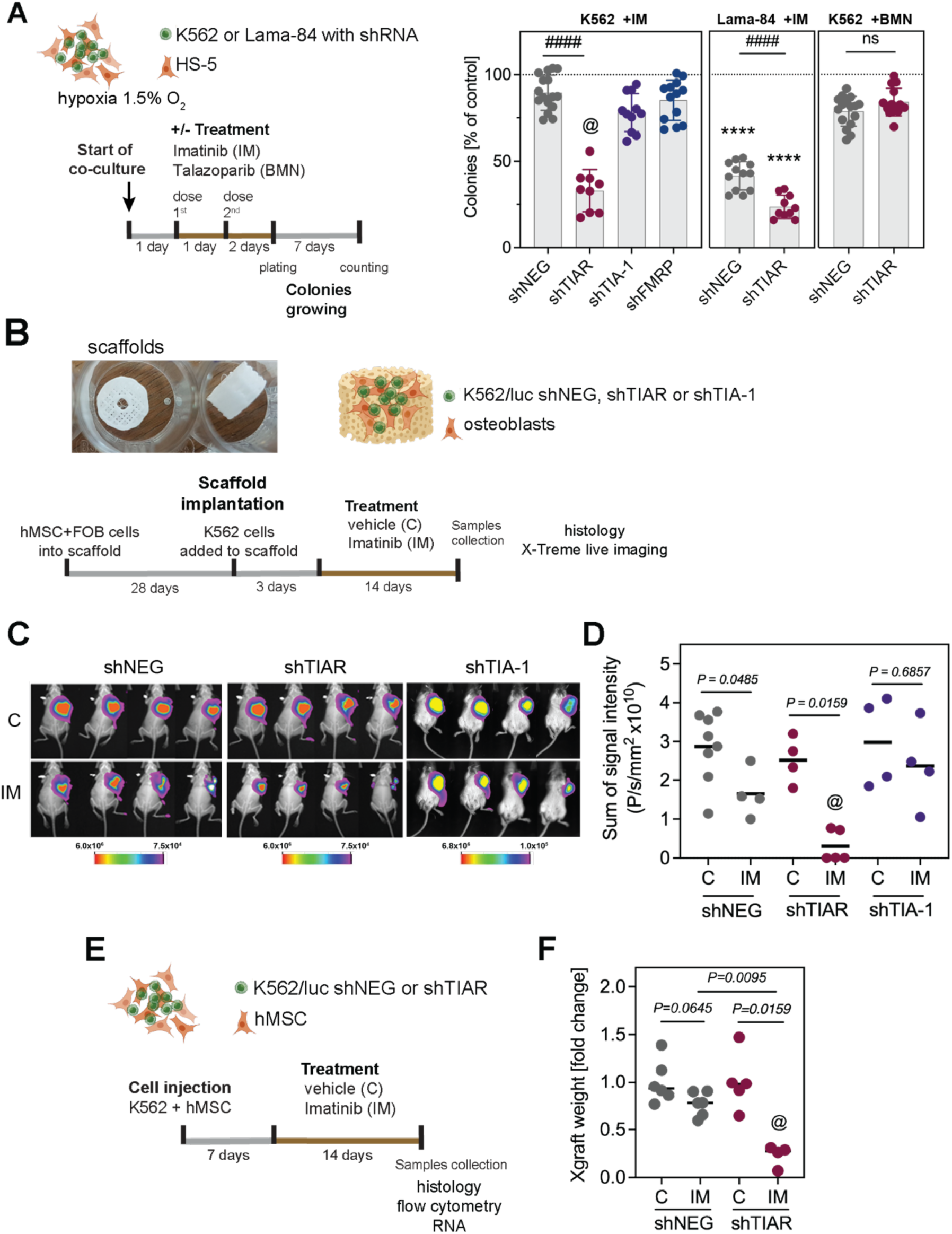
Silencing of TIAR sensitizes leukemia cells to imatinib treatment in different setups modelling human bone marrow stroma *ex vivo* and *in vivo*. **(A)** Colony formation by K562 cells expressing shRNA non-targeting (shNEG) or targeting mRNA of *TIAL1* (shTIAR), *TIA1* (shTIA-1), or *FMR1* (shFMRP) that were collected from *ex vivo* hypoxic (1.5% O_2_) co-culture with HS-5 bone marrow stromal fibroblasts and treated with 1 μM imatinib (IM) or 50nM Talazoparib (BMN) added in two doses following the experimental scheme (left panel). Number of colonies in each of the 3 technical replicates from 3-4 independent biological experiments presented as % change relative to the untreated cells (dashed black line) set as 100 %. Student’s two-tailed t-test was used to compare two samples marked by the black line; #### or **** - p<0.0001, ns - p > 0.05. **(B)** Scheme explaining the experimental setup based on the subcutaneous implantation in mice of 3D printed scaffolds (photo taken with a Samsung mobile phone camera) seeded with human cells differentiated into osteoblasts and K562/luc cells with shNEG, shTIAR or shTIA-1, followed by IM treatment for 14 days. **(C)** Bioluminescence signal monitored in mice after 2 weeks of IM or vehicle treatment (timeline explained in **(B)**) following luciferin injection, collected in the Burker’s Xtreme In-Vivo chamber for 30 sec, and overlaid on the mouse X-ray image. Scale presents the signal intensity of the color coding. **(D)** Sum of the signal intensity (P) collected per second (s) and area (mm^2^) for each mouse analyzed (single dot) is presented. **(E)** Scheme explaining experimental steps of xenograft formation by K562/luc (expressing shNEG or shTIAR, Firefly luciferase, and GFP) mixed with human primary bone marrow mesenchymal stem cells (hMSC) subcutaneously injected in mice 7 days before initiation of treatment with IM for the following 14 days. **(F)** Weight of each xenograft isolated from mice (single dot) formed by K562/luc cells with shNEG or shTIAR (as in **(E)**) presented as fold change of the weight mean value of xenografts from mice treated with vehicle; mean value indicated with the black line. The nonparametric two-tailed Mann-Whitney test was used for comparisons indicated by black lines underneath the exact significance (*P*) values are presented in the plot. **(A,D,F)** Two-way Anova was used to compare shTIAR upon IM versus other variants; @ - P < 0.002.

### TIAR is necessary for CML cell persistence within the bone marrow stroma niche upon imatinib treatment

Cell adaptation to the hypoxic BM microenvironment necessitates extensive proteome metabolic rewiring to sustain cell survival. Transcriptomic data from the xenograft model point to translation as a process significantly modulated by hMSCs in leukemia cells (Fig. S3A). Our previous work has shown that the BM microenvironment profoundly influences translation regulation and has identified a critical role for RNA-binding proteins in this process in CML cells (*10, 34, 35, 46*). Notably, in response to translation inhibition TIAR knockdown reduced CML cell survival in experiments performed under hypoxic conditions but not under atmospheric O_2_ (normoxic) (*10*). We observed that in CML patients resistant to TKIs, the level of TIAR expression was increased (Fig. S5A). Building on these findings, we investigated the effect of TIAR silencing on the relapse potential of CML cells after IM treatment under hypoxic co-culture conditions (Fig. 4A). Using previously described cellular models (*10*), we tested the impact of several RBPs (Fig. S5B). In K562 cells, shTIAR reduced colony formation by 57 ± 4%, p<0.0001, with no significant effect observed in shNEG controls. Silencing of other RBPs that respond to the integrated stress response activation, Tia-1 or FMRP, yielded no substantial effect and reduced colony numbers by 12 ± 4% (p=0.0101) and 4 ± 4% (p=0.2955), respectively. In another CML cell line, Lama-84, IM treatment alone reduced colony formation by shNEG cells by 59 ± 2% (p<0.0001), and shTIAR further decreased this by an additional 18 ± 3% (p<0.0001). Importantly, the effect of TIAR knockdown was specific to IM, with no significant impact on the efficacy of the PARP1 inhibitor Talazoparib (BMN). To further characterize the cellular basis of these effects, we assessed the impact of shTIAR on cell viability under hypoxic co-culture conditions. IM treatment for 48h increased the number of cells with fragmented DNA (subG1 fraction) (Fig. S6A), shifted cells into the G1/G0 cell cycle with a concomitant reduction in G2/M fraction (Fig. S6B), and promoted apoptosis (Fig. S6C). TIAR knockdown selectively reduced cell number in the subG1 fraction without significantly affecting cell cycle progression or cellular apoptotic signaling (Fig. S6D).

To evaluate the role of TIAR in leukemia cell survival *in vivo,* we employed two complementary experimental approaches (Fig. 4B-F). First, to recapitulate the three-dimensional architecture of the BM, we used 3D-printed scaffolds seeded with hMSC-derived osteoblasts, into which K562/luc cells were introduced prior to subcutaneous implantation in mice (Fig. 4B). Following 2 weeks of IM treatment, leukemia cell expansion within scaffold was assessed by bioluminescence imaging of luciferase activity (Fig. 4C-D). TIAR silencing produced the greatest reduction in bioluminescence signal among all conditions tested, including shNEG and shTIA-1, significantly decreasing the IM-to vehicle-treated signal by 2.2 ± 0.3 in shTIAR mice (Fig. 4D). No morphological differences were detected in scaffold cryosection staining across conditions (Fig. S7A). Of note, IM treatment alone significantly reduced the shNEG bioluminescence signal in scaffolds by 1.2 ± 0.5 relative to vehicle-treated controls (Fig. 4D). To corroborate these findings, K562/luc cells were co-implanted with undifferentiated hMSC to generate subcutaneous xenografts (Fig. 4E). Following 2 weeks of IM treatment, we checked the xenograft weight (Fig. 4F). IM treatment had no significant effect on shNEG xenograft weight, indicating that this stromal configuration conferred more robust pro-survival support to leukemia cells. Strikingly, TIAR silencing significantly reduced xenograft weight by 53 ± 0.1% upon IM treatment (Fig. 4F), without affecting spleen size (Fig. S7B).

Together, results across multiple experimental systems demonstrate that TIAR knockdown selectively attenuates the survival potential of IM-treated CML cells, overcoming the pro-survival support presented by the human BM stromal microenvironment, independently of effects on apoptosis or cell cycle progression.

### TIAR is required for IM-induced transcriptional and alternative splicing reprogramming in CML cells

We determined that IM-induced transcriptional changes in xenograft-derived leukemia cells are largely dependent on TIAR. RNA sequencing of FACS-isolated cells from xenografts (as characterized in Figure 4E-F) revealed that nearly 68% of genes upregulated by IM treatment in shNEG cells were downregulated upon TIAR silencing, while 86% of IM-downregulated genes were reciprocally upregulated in shTIAR cells (Fig. 5A). Functional annotation of these inversely regulated genes encoded proteins involved in transcriptional regulation, transmembrane transport, and metabolism (Fig. 5B). This pattern of pathway enrichment was recapitulated in an *ex vivo* co-culture system, where the most prominently affected biological processes were cell adhesion and ion transmembrane transport (Fig. S8).

**Figure 5.**
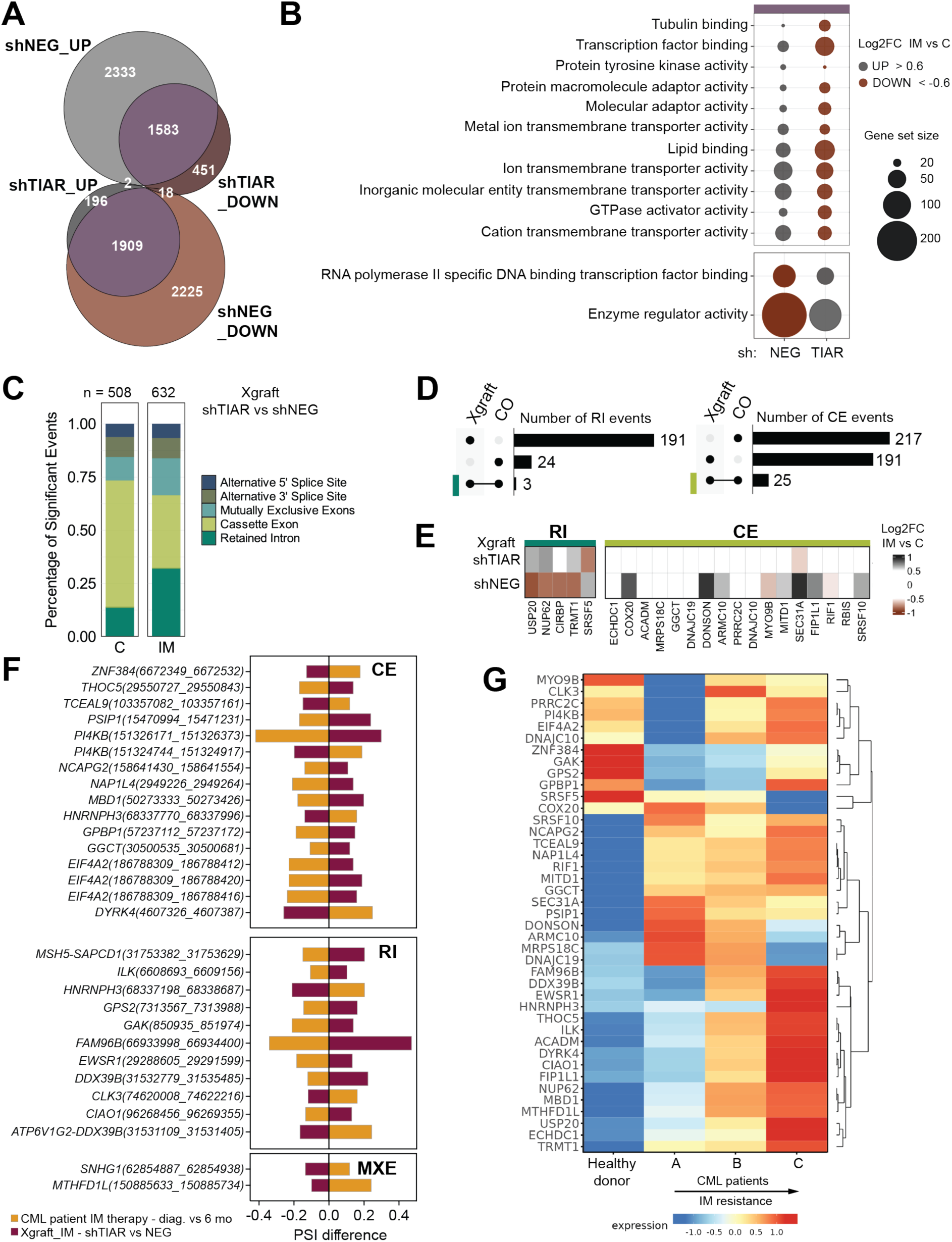
Silencing of TIAR reverses the changes in gene expression induced by imatinib treatment in leukemia cells supported by the bone marrow stroma. **(A)** Comparison of gene expression in K562/luc cells with shNEG or shTIAR, FACS-sorted from xenografts (Xgraft), with level upon 2 weeks of IM versus vehicle (C) treatment changed with log2 value of fold change (Log2FC) ≥ 0.6 (UP) or ≤-0.6 (DOWN); reversed regulation in shTIAR cells in purple. **(B)** Gene Ontology Molecular Function terms identified by the GSEA of genes that upon IM vs C in Xgraft of shNEG are reversely regulated in shTIAR (shaded in purple). In grey circles - UP (Log2FC > 0.6) n = 1993, brown circles DOWN (Log2FC <-0.6) n = 1756. Selected terms with NES_abs_ ≥ 1.45, p-value ≤ 0.05, and the number of genes in the sample annotated per term (size) ≥ 10; n-number of genes. **(C)** Proportion of alternative splicing events (AS) changed upon shTIAR versus shNEG in cells from xenografts treated with vehicle (C) or imatinib (IM) for 2 weeks. Significant events from rMATS analysis requiring at least 20 reads/event, absolute change in PSI (Percent Spliced In) > 0.1 with FDR ≤ 0.05; total number in 3 experiments (n) above the bar. **(D)** Comparison of intron retention (RI; left panel) and cassette exon (CE; right panel) AS with significant PSI changed in shTIAR versus shNEG in cells from Xgraft or CO treated with IM. **(E)** Change in expression level of genes in Xgraft IM versus vehicle-treated for a subset of genes with significant changes shTIAR vs shNEG in RI (left panel) or SE (right panel) in cells from Xgraft and CO treated with IM; Log2FC ≤-0.6 in brown, ≥ 0.6 in grey. **(F)** Difference in PSI value of CE, RI, and mutually exclusive exons (MXE) AS in: moro – shTIAR versus shNEG xenograft cells from mice treated with IM (Xgraft_IM); yellow – CD34^+^ enriched cells from bone marrow biopsies of a CML patient collected at 6-month IM therapy versus diagnosis GSE310243 (see in Figure 3B). Included only with PSI > 0.1 and with FDR ≤ 0.05. **(G)** Expression level of genes with changes in AS (selected in **(E)** and **(F)**) analyzed at the scdbm (*45*) for the CD34^+^ cells subtype. Patient classification in Figure 3D.

Beyond transcript abundance, altered transcript processing expands the isoform repertoire in cancer cells, enabling leukemia cell therapy resistance (*47*). IM treatment induced changes in alternative splicing (AS), involving cassette exons (CE) inclusion, mutually exclusive exons (MXE) usage, and retention of introns (RI), detected in both co-culture and xenograft systems (Fig. S9A). Notably, the greater transcriptome-wide changes observed in xenografts relative to co-cultures (Fig. S10) likely reflect the longer duration of IM exposure and so more extensive AS remodeling. TIAR silencing reduced variability in AS events in both experimental systems and reduced the number of AS changes upon IM treatment (Fig. S9A). Knockdown of TIAR influenced CE, RI and MXE (Fig. 5C), affecting a greater number of events under control conditions than upon IM treatment (Fig. S9B). Most of the splicing changes caused by shTIAR were condition-dependent (Fig. S9C), and under IM treatment preferentially involved increased intron or cassette exon inclusion rather than exclusion (Fig. S9D). AS changes were accompanied by variability in overall intron splicing efficiency, which was more pronounced in cells derived from xenograft (Fig. S9E) than from co-culture (Fig. S9F). A subset of RI and CE events affected by shTIAR under IM treatment was reproducible across experimental systems (Fig. 5D) and was accompanied by concordant changes in the expression of the corresponding genes (Fig. 5E). Furthermore, comparison with biopsy data from an individual IM-treated CML patient demonstrated that TIAR targeting reversed the RNA-processing effects of IM therapy observed in the patient (Fig. 5F, S11). Strikingly, most of the transcripts exhibiting modified exon or intron inclusion were upregulated in CD34-positive BM cells from CML patients with IM treatment failure or resistance (Fig. 5G).

Collectively, these findings point to TIAR as a critical regulator of IM-induced AS reprogramming, implicating alternative splicing as a key component of the transcriptomic adaptation that enables CML cell persistence in the BM under therapeutic pressure.

### TIAR modulates alternative splicing and nascent protein synthesis upon IM treatment

The nascent proteome reflects the selectivity of mRNA translation modulated by RBPs such as TIAR and FMRP, which is microenvironment-dependent in CML cells (*10*). Therefore, we characterized IM-induced changes in the nascent proteome of CML cells under hypoxic co-culture conditions (Fig. 6A). To this end, we employed QuaNCAT, an approach combining biorthogonal noncanonical amino acid tagging (BONCAT) with stable isotope labeling by amino acids in cell culture (SILAC) using ‘heavy’ or ‘medium’ Arg and Lys, to quantify nascent protein levels in shTIAR versus shNEG cells, as previously described (*10*). Among nascent proteins identified, 162 proteins were significantly changed (p < 0.05, absolute fold change > 50%). Simultaneously, 915 mRNAs were differentially expressed, and only 14 mirrored their corresponding protein change (Fig. 6B). Among the 66 upregulated proteins, the most enriched functional categories included RNA-binding proteins, hemoglobin-associated proteins, oxidoreductases and peroxidases (Fig. 6C, upper panel). The 96 downregulated proteins were predominantly associated with cell adhesion, lipid metabolism, transmembrane transporters, and ubiquitination (Fig. 6C, lower panel).

**Figure 6.**
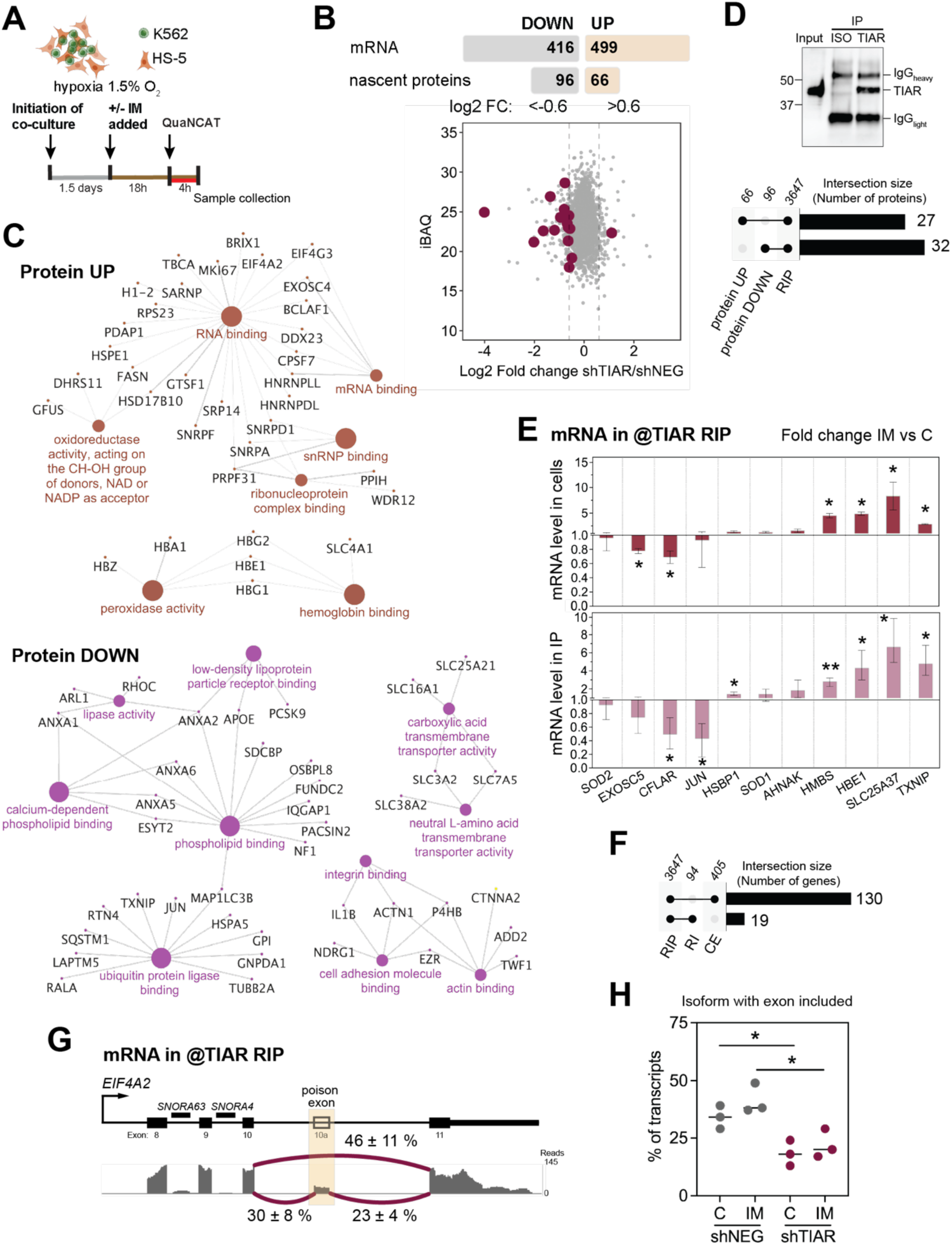
TIAR modulates protein synthesis and alternative splicing of RNA targets upon imatinib treatment. **(A)** Scheme explaining experimental setup to determine proteomic changes using the quantitative BONCAT (QuaNCAT). K562 cells expressing shNEG or shTIAR growing in co-culture with HS-5 cells under hypoxia (1.5% O_2_) for 1.5 days were treated with imatinib (IM) for 18h before the bioorthogonal noncanonical amino acid tagging (BONCAT) of nascent proteins synthesized in cells for 4h. **(B) Upper panel** - number of genes and nascent proteins showing increased (UP; log₂ fold change [Log₂FC] ≥ 0.6) or decreased (DOWN; Log₂FC ≤ −0.6) abundance in IM-treated shTIAR cells compared with shNEG cells, with a significance threshold of p ≤ 0.05. In total, 2186 nascent proteins were quantified in the BONCAT experiment. **Lower panel** - scatter plot showing intensity-based absolute quantification (iBAQ), used as an estimate of relative protein molar abundance, plotted against Log₂FC values for BONCAT-identified proteins in shTIAR versus shNEG cells. Purple dots indicate hits with significant changes at both the protein and mRNA levels. Dashed vertical lines mark the Log₂FC thresholds of −0.6 and 0.6. **(C)** Functional annotation Gene Ontology Molecular Function enrichment analysis of UP (brown) or DOWN (purple) proteins with ClueGO/CytoScape, displaying proteins annotated with the term; only one side enriched terms with FDR < 0.05. **(D) Lower panel** - number of proteins that upon shTIAR are UP and DOWN regulated, for which mRNA was detected in the RNA immunoprecipitated (IP) in TIAR protein complexes (RIP), enriched in samples obtained with anti-TIAR antibody versus the same isotype non-binding antibody (ISO) (n=3647). **Upper panel** - example Image of Western blotting analysis of IP, with a sample of cell lysate used for IP loaded for reference (input). **(E)** Changes in the mRNA level, determined by real-time PCR and quantified using the ddCT method, expressed as log2 of change between IM-treated versus untreated cells, detected in samples from whole cells (upper panel) and anti-TIAR RIP (lower panel). Mean of 3 independent experiments with ± range is presented. Student’s t-test two-way was used to compare the difference between IM to C; * p ≤ 0.05, ** p≤0.005. **(F)** Number of genes identified in anti-TIAR RIP that show significant changes in intron retention (RI) or cassette exon alternative splicing (CE) in shTIAR versus shNEG alternative splicing analysis of RNA from IM-treated cells. Number of genes in the intersection provided above the bar; comparisons indicated by the black dot. **(G)** Sashimi plot (middle panel) demonstrating splicing of *EIF4A2* mRNA within the region encompassing exons 8-11 and 3’UTR in RNA from anti-TIAR RIP. Alternative splice site usage marked by the purple line, and the percent usage ± ME (n=3) in numbers by the lines, reads coverage from 0-145 shown in grey, alternatively spliced exon marked by shaded yellow. Gene region scheme in the top. **(H)** Percent transcripts with TIAR-dependent alternative exon inclusion in K562 cells from xenografts treated with IM. Student’s t-test was used to compare results from three experiments (each dot) for the mean value marked with a thick black line; * p = 0.021.

Analysis of UV cross-linking RNA immunoprecipitated (UV-RIP) with TIAR from hypoxia co-cultured CML cells indicated that the synthesis of 59 of the significantly changed proteins (∼ 36%) may be directly regulated through TIAR-mediated post-transcriptional modulation (Fig. 6D). This corroborated with detection of IM-induced changes in binding of the corresponding mRNAs by real-time PCR analysis of UV-RIP fractions (Fig. 6E). Furthermore, nearly 30% of transcripts exhibiting significant AS changes upon TIAR knockdown were directly targeted by TIAR, accounting for 20% of RI and 32% of CE events (Fig. 6F, S12, Table S1). Similarly, in IM-treated shTIAR cells, changes (UP or DOWN) in protein synthesis coincided with changes in mRNA level in only 16 cases (Fig. S13A). Approximately 20% of TIAR-bound mRNAs were downregulated and almost 10% upregulated (Fig. S13B), and shTIAR-dependent AS changes overlapped with nascent protein synthesis alterations in 21 cases (Fig. S13C). Among the upregulated proteins (Fig. 6C) was eukaryotic initiation factor 4 A2 (EIF4A2). In experiments with CML cells performed under normoxic conditions, we have previously found that 18 hours of IM treatment reduced the inclusion of a poison cassette exon in *EIF4A2* transcripts (*27*), a change recognized to induce nonsense-mediated decay (*48*). Here, about half of the TIAR-immunoprecipitated *EIF4A2* transcripts exhibited this exon skipping (Fig. 6G). Silencing of TIAR significantly reduced poison cassette exon inclusion in *EIF4A2* transcripts isolated from xenograft-derived leukemia cells, independently of IM treatment (Fig. 6H, S12). This effect phenocopies the IM-induced splicing change observed under normoxia (*27*), in which CML cells retain IM sensitivity (*21*). This suggests that TIAR-dependent regulation of EIF4A2 splicing may represent an adaptive mechanism governed by oxygen availability.

Integrated analysis of the transcriptome, nascent proteome, and TIAR complex-associated RNA revealed that TIAR-modulated targets converge on the intracellular antioxidant system and redox regulation, coinciding with erythroid differentiation.

### Imatinib-induced erythroid differentiation is attenuated by TIAR silencing

IM treatment is known to induce erythroid differentiation in CML cells, driving expression of hemoglobin genes in K562 cells (*49–51*) and accompanied by transcriptional readthrough, as we reported recently (*27*). Erythroid-biased progenitors dominated in the bone marrow of IM-responding patients and were more sensitive to TKIs (*45*). Consistently, in CD34-positive cells from CML patient BM (*45*), IM therapy upregulated expression of genes encoding proteins with essential roles in erythropoiesis that were also affected by shTIAR, including transferrin receptor 2 (*TFR2*), GATA-binding protein 1 (*GATA1*), fatty acid translocase (*CD36*), hemoglobin subunit epsilon 1 (*HBE1*), hemoglobin subunit gamma 2 (*HBG2*), and ferritin light chain (*FTL*) (Fig. 7A). Western blot analysis of cell lysates corroborated the shTIAR induced changes in nascent protein levels observed upon IM treatment (Fig. 6C), except for thioredoxin-interacting protein (TXNIP) (Fig. 7B), which could result from increased protein stability mediated by thioredoxin activity (*52*). Dysregulated levels of proteins involved in lipid peroxidation, oxidoreduction, and transmembrane transport were associated with reduced mitochondrial membrane potential (Fig. 7C) and cellular reduction potential (Fig. S14A), accompanied by accumulation of lipid hydroperoxides (LOOH) (Fig. 7D). Simultaneously, hydrogen peroxide levels were unaffected (Fig. S14B), consistent with the lack of TIAR knockdown influence on nucleic acid damage (Fig. S14C).

**Figure 7.**
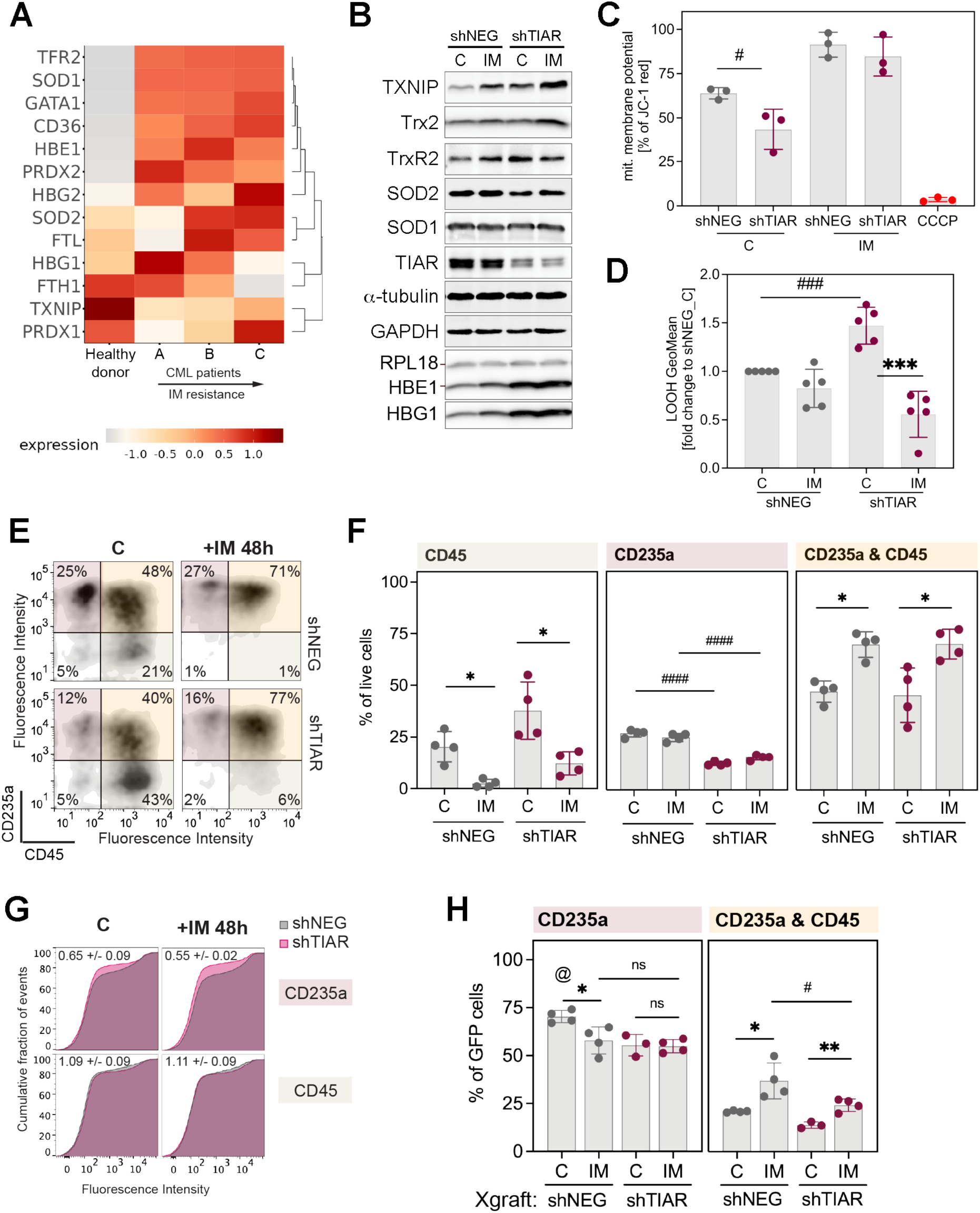
Silencing of TIAR affects the cellular redox state and reduces the number of erythroid-differentiated cells. **(A)** Expression level of genes analyzed at the scdbm (*45*) for CD34^+^ cells subtype; patient classification explained in Figure 3D. **(B-H)** Impact of shTIAR compared to shNEG analyzed in K562 cells that were co-cultured with HS-5 cells under hypoxia (1.5%O_2_) 2 days before initiation of 18h culture with imatinib (IM) treatment or without (C). **(B)** Protein level in whole cell extracts analyzed by Western blot, representative images of immunoblots (n=3) presented. **(C)** Mitochondrial (mit.) membrane potential measured using JC-1 probe; values for signal from probe aggregates (red) in polarized mitochondria expressed as % of signal from the probe in the cells (n=3), and carbonyl cyanide 3-chlorophenylhydrazone (CCCP) used as a control. **(C,D,F,H)** Bars represent the mean of values from independent experiments (indicated by dots) with ± SD. Student’s t-test was used to determine the significance of the difference shTIAR vs shNEG (#) and IM vs C (*); * - p ≤ 0.05, ** p ≤ 0.01, *** p ≤ 0.005, **** p ≤ 0.001 **(D)** Lipid peroxidation measured with click-it chemistry by flow cytometry. Fluorescence intensity GeoMean expressed as fold change of value in the untreated shNEG cells (n=5). **(E-H)** Presentation on the cluster of cell differentiation (CD) surface protein markers CD45 and CD235a on K562 cells with shNEG or shTIAR isolated from co-culture established in hypoxia (1.5%O_2_) a day before initiation of treatment with IM for 48h **(E-G)** or FACS-sorted K562/luc GFP positive cells from xenografts **(H)**. **(E-F,H)** Percentage of parental live cell subpopulations that are: double positive for CD45 and CD235a (CD235a&CD45), positive only for CD45, or only for CD235a, or negative for both CD markers. Representative scatter plots in **(E)**, summary of independent co-culture experiments (n=4) in **(F)**. **(G)** Fluorescence intensity of surface CD36 protein staining in the fraction of cells positive for CD235a or CD45. Numbers correspond to fold change in shTIAR to shNEG ± ME (n=2). **(H)** Percent GFP and hCD45-positive cells from different xenografts (n=4) that are positive for CD235a. Two-way ANOVA test was used to for comparison of shNEG_C to other variants (@), and p=0.0046.

The concurrent occurrence of erythroid differentiation and LOOH accumulation suggested that TIAR silencing promotes ferroptosis, which is an iron-dependent form of regulated cell death. In support of this, TIAR knockdown in the pool of viable cells significantly reduced the proportion of CD45-negative/glycophorin A (CD235a)-positive cells (Fig. 7E-F), thus displaying a surface phenotype characteristic of erythropoietic-biased cells. Besides, within the CD235a-but not CD45-positive cell population, shTIAR reduced the population presenting surface fatty acid translocase (CD36), a transmembrane receptor mediating lipid uptake (Fig. 7G). Markedly, xenograft-derived cells exhibited the same phenotypic characteristics (Fig. 7H), reinforcing the CML relevance of these findings.

## DISCUSSION

To investigate CML cell response to IM treatment within a human BM stromal microenvironment, we employed three complementary in vivo approaches: (i) formation of hMSC-derived ossicles colonized by CML cells, (ii) 3D-printed scaffolds seeded with hMSC-differentiated osteoblasts into which CML cells were introduced before implantation, and (iii) subcutaneous xenotransplantation of CML cells co-implanted with undifferentiated hMSCs. Across all three systems, CML cells persisted following 2 weeks of IM treatment within the BM stromal microenvironment. Of the three approaches, ossicles proved to be the least robust and the most time-consuming. Scaffolds, while oriented at recapitulating the bone-oriented endosteal niche, precluded RNA-seq analysis of recovered cells. Interestingly, IM exerted a more pronounced effect on CML cell survival in this system. This could be attributed to better vascularization resulting in a less hypoxic local environment, or to an insufficient abundance of hMSCs or other hMSC-derived cell types such as adipocytes. Formation of xenografts enabled control of the number of transplanted cells and efficient recovery of IM-resistant CML cells at the end of the experiment. Here, to characterize BM stromal niche-dependent changes in RNA processing, we integrated xenograft model data with published datasets obtained from analysis of BM biopsies. The xenograft model recapitulated a subset of gene expression and AS alterations also detected in CML cells from BM patient samples, demonstrating translational validity of the system.

We proved that alternative splicing remodeling and transcriptomic reprogramming are integral components of the mechanism underlying CML cell survival within the BM stromal microenvironment and are modulated by stromal cell interaction. Remarkably, changes in the synthesis of selected proteins involved in IM-altered processes identified in long-term treated CML cells *in vivo* were detectable already within 18 hours of treatment initiation in the *ex vivo* leukemia cells co-cultured with BM fibroblasts in hypoxia. This corroborates our previous finding that a subset of transcriptomic changes detected in CML cells surviving long-term IM therapy are initiated shortly after treatment onset (*27*), and now extends this observation to the BM stromal niche.

We identified TIAR as a critical player supporting CML cell survival within the BM microenvironment during IM treatment. Studies conducted in the distinct biological context of epithelial cancers have suggested a tumor-suppressive role for TIAR (*53, 54*), reasoned from observations that reduced TIAR expression enhanced cell proliferation, while its overexpression suppressed xenograft growth *in vivo* and modulated the expression of regulators of the cytoskeletal organization, cell adhesion, and cell cycle progression (*37, 38, 54, 55*). In the present study, TIAR silencing under hypoxic co-culture conditions similarly altered the expression of factors involved in these processes, but no difference in leukemia cell proliferation was observed upon TIAR knockdown in the *ex vivo* hypoxic co-culture model, neither in control conditions (*10*) nor upon IM treatment. This discrepancy could result from the anchorage-independent nature of K562 cell cycling in culture. Apart from that, we have previously evidenced that the TIAR function is microenvironment-dependent, particularly in the context of treatment response (*10*). In leukemia, hypoxic BM-enforced dormancy is the primary determinant of therapy survival, such that disruption of cell cycle arrest maintenance would be expected to reduce, rather than enhance, CML cell persistence. As cells with TIAR depletion display increased Ki-67 (*MKI67*) nascent protein synthesis under IM treatment, a canonical marker of active cell division, it is plausible that TIAR knockdown could impair the acquisition of quiescence, thereby undermining a mechanism essential for CML cell survival under therapeutic pressure.

Silencing of TIAR could attenuate CML cell persistence in the BM by affecting metabolic adjustments induced by IM treatment. Many of the metabolically significant changes associated with IM survival are reversed upon TIAR silencing. The impact of TIAR depletion on gene expression likely reflects disruption of direct RBP-RNA interactions that control both transcript splicing and translation. We identified that TIAR silencing altered AS or protein synthesis of several metabolic enzymes. For instance, upon IM therapy shTIAR induced inclusion of cassette exon, which in CML patient samples is more skipped in gamma-glutamylcyclotransferase (GGCT), an enzyme recently implicated in the coordination of glucose metabolism and cell proliferation in tumor cells (*56*). Moreover, TIAR silencing affected the protein synthesis of Trx-2 or SOD2, enzymes critical for reactive oxygen species scavenging, as well as fatty acid synthase (FASN), dehydrogenase/reductase 11 (DHRS11) and pyruvate kinase M (PKM), all involved in glucose and lipid metabolism. In addition to metabolic enzymes and proteins of the antioxidant system, the dysregulated level of SLC transporters upon TIAR depletion likely contribute to the observed impairment of cellular redox potential.

TIAR coordination of RNA processing and translation could support metabolic rewiring, which is well recognized as a driver of therapy resistance in leukemia. We found changes in PKM splicing, and this shift in the enzyme isoform is linked to therapy resistance in acute lymphoblastic and myeloid leukemia (*57, 58*). Quiescent cells rely on fatty acid oxidation (FAO) for ATP generation (*59*), a process modulated by BM adipocytes (*60*) and representing an exploitable therapeutic vulnerability in leukemia (*61–63*). Moreover, the FAO-driven metabolic rewiring enforces cell quiescence in hematopoietic cells (*64*). The observed here accumulation of lipid peroxides, can be fueled by elevated FAO and CD36-mediated fatty acid uptake and is a well-documented trigger of ferroptosis across multiple cancer types (*65, 66*). Our analysis revealed that increased hemoglobin synthesis was accompanied by a reduced number of viable erythroid-differentiated cells. Moreover, we observed that the viable CD235a-positive population with surface CD36 was reduced upon TIAR silencing. Taken together, these findings indicate that TIAR silencing in IM-treated CML cells under hypoxic conditions promotes cell death by ferroptosis, thereby potentiating the therapeutic efficacy of IM.

In conclusion, we demonstrated that TIAR represents a critical node within the regulatory network coordinating RNA processing and cell redox state, detrimental to metabolic rewiring that sustains CML cell persistence in the BM microenvironment under IM treatment. By employing experimental systems that recapitulate key aspects of therapy resistance, we uncovered a previously unrecognized cell survival strategy based on coordinated stromal niche-driven changes in RNA processing and metabolism. This reveals a new vulnerability of CML cells that could be exploited to improve therapeutic outcomes.

## MATERIALS AND METHODS

### Cell culture

Human chronic myeloid leukemia cell line K562 (#CCL-243) and bone marrow stroma fibroblast cell line HS-5 (#CRL-11882) were obtained from American Type Culture Collection (USA); human chronic myeloid leukemia cell line LAMA-84 (#ACC 168) was from DSMZ. Cells were cultured in RPMI-1640 (Biowest, France, #L0490) supplemented with 100 U/ml penicillin, 100 μg/ml streptomycin (Biowest, #L0022), 2 mM L-Glutamine (Biowest, #X0550) and 10% FBS (vol/vol; Biowest, #S1810) as previously described (*10*). For 72 h co-culture, cells were seeded at 1:1 leukemia-to-stroma ratio with HS-5 cells. Cell cultures were grown in a CO_2_ incubator at 37°C, 5% CO_2,_ and atmospheric O_2_. The co-cultures were kept in the InVivo 400 hypoxia workstation (Baker Ruskinn, UK) at 37°C, 5% CO_2,_ and 1.5% O_2_. Handling steps, such as the exchange of medium or the addition of treatment, were also performed within the workstation.

### Human primary cells

Human Bone Marrow-Derived Mesenchymal Stem Cells (MSC) used for *in vivo* experiments were from ATCC (#ATCC-PCS-500-012) and were cultured initially in the MSC basal medium (#ATCC-PCS-500-030) with supplement (#ATCC-PCS-500-041). From the third passage, MSC cells were cultured in DMEM (Sigma-Aldrich, #D5546) supplemented with 100 U/ml penicillin, 100 μg/ml streptomycin, 2 mM L-Glutamine, and 10% FBS. The primary cells from CML-CP patients (2 donors) were also used in other projects (*22, 44*) and provided by Prof. T. Skorski. The enrichment for CD34-positive cells from the mononuclear fraction and *ex vivo* expansion preceding mouse injection were performed as previously described (*9*). The samples were subjected to negative selection of progenitor cells (Lin-) followed by enrichment for CD34-positive cells by a magnet separation after immunostaining with antibodies conjugated to magnetic beads (StemCell Technologies #17936 and #17856). The primary Lin-CD34+ cells were expanded in medium StemPro-34 SFM (ThermoFisher Scientific #10639011) supplemented with 2 mM L-Glutamine and growth factors from Peprotech: 100 ng/mL SCF (#300-07-10UG), 100 ng/mL Flt3 ligand (#300-19-10UG), 20 ng/mL IL-3 (#200-03-10UG), 20 ng/mL IL-6 (#200-06-20uG).

### Lentiviral transduction

For stable gene knockdown, the MISSION® shRNA lentiviral particles from Merck/Sigma Aldrich (St. Louis, MO, USA) were used. Detailed protocol of K562 cell transduction with MISSION® shRNA Lentiviral Transduction Particles for TIAR knockdown (shTIAR) or scrambled control (shNEG) has been described previously (*10*). Following 2 weeks of selection in 1 µg/ml puromycin for stably transduced cells, the cells were subjected to a second lentiviral transduction with plasmid pLenti7.3-redluc (Invitrogen #V53406) to generate cells stably expressing GFP and Firefly luciferase (K562/luc). GFP-positive cells were enriched to 95% by flow cytometry cell sorting with BD FACS Aria (RRID:SCR_018934). The level of gene silencing was verified with RT-qPCR and Western Blot analysis.

### *In vitro* drug treatment

Imatinib (IM; a generous gift from the Pharmaceutical Institute in Warsaw) was used to inhibit BCR-Abl1 oncoprotein activity. To this end, the 10 mM stock of IM mesylate solution in dimethyl sulfoxide (DMSO; SigmaAldrich #4540) was diluted in RPMI-1640 to 1 mM and then added to the cell culture at a final concentration of 1 μM.

### Colony-forming assay

Colony-forming assay was performed as previously described (*9, 10*). Briefly, HS-5 cells were seeded 48 h before the addition of K562. After 24h, the first dose of IM at 1 μM or BMN at 50nM final concentration was added to the co-culture, followed by a second dose after 24h. After another 48h, the cells were collected and resuspended in methylcellulose-based medium (StemCell, #H4230) supplemented with 10% FBS. The colonies formed by the cells that survived the treatment were counted after 7 days.

### Cell lysis and Western Blotting

Total cell lysates were prepared in hot SDS lysis buffer following a protocol described in detail previously (*10*). Briefly, after washing with PBS, cells were resuspended in the lysis buffer (50 mM Tris-HCl pH 6.8, 10% glycerol, 2% SDS) and incubated for 8 min at 95 °C. The same amount of protein μg was subjected to SDS-PAGE followed by Western Blotting analysis. Antibodies used to develop the Western blots were: α-tubulin (Calbiochem, #CP06-100UG), TIAR (Bethyl, #A303-613A), HBE1 (Proteintech, #12361-1), HBG1 (Proteintech, #25728-1), RPL18 (Proteintech, #17029-1), TXNIP (Proteintech #18243-1), Trx2 (SantaCruz Biotech., #sc-133201), TrxR2 (SantaCruz Biotech., #sc-166259), SOD1 (SantaCruz Biotech., #sc-101523), SOD2 (SantaCruz Biotech., #sc-137254), GAPDH (Bethyl, #A300-639A), and secondary antibodies conjugated to horseradish peroxidase: goat anti-mouse (Dako, #P0447) and goat anti-rabbit (Dako, #P0448). Western Blots were imaged and analyzed using ChemiDoc MP Imaging system (BioRad, RRID:SCR_019037).

### Antibody Arrays

Apoptotic signaling was checked in lysates with 1mg of protein (10x diluted) of K562 shNEG and shTIAR cells treated with IM for 24h. To this end, the Human Apoptosis Signaling Array was used (RayBiotech #AAH-APO-1) following the manufacturer’s protocol.

### QuaNCAT sample preparation

Cell labeling, sample preparation for the LC-MS/MS, and data analysis were performed as described in detail previously (*10*). To this end, K562 cells were co-cultured with HS-5 cells under hypoxia for 48h before IM was added for 18h. During the final 4 h, newly synthesized proteins were metabolically labeled by incorporation of the methionine analog azidohomoalanine together with stable isotope-labeled arginine and lysine. Raw files corresponding to 3 replicate samples obtained from shTIAR (Heavy, H) and shNEG (Medium, M) were processed together. For analysis of data, protein groups for which all the detected Log2 H/M values of 3 experiments were > 0, and at least two Log2 H/M values were > 0.6 were classified as upregulated in shTIAR (UP). Protein groups for which all the detected Log2 H/M values were < 0, and at least two Log2 H/M values were <-0.6 were classified as downregulated in shTIAR (DOWN). The protein network enrichment analysis of GOMF (updated Nov 2024) was performed using the ClueGO plug-in at CytoScape (v3.10.3) (RRID: SCR_003032), with a list of identified proteins in either condition as a reference, displaying only pathways with significant enrichment (Right-sided hypergeometric test) corrected with Benjamini-Hochberg FDR set to ≤ 0.05.

### RNA immunoprecipitation (UV-RIP)

Protein-RNA interactions were crosslinked by subjecting the cells to two rounds of illumination at 254nm (200mMJ/cm2) for 1min in the Spectrolinker XL-1500 UV crosslinker (Spectronics). To this end, cells were suspended in 10ml of ice-cold PBS and placed in 100mm plates on a tray filled with ice. After two washes in ice-cold PBS, the cell pellet was resuspended in a lysis buffer (10 mM HEPES KOH pH 7.2, 150 mM KCl, 5 mM MgCl_2_, 0.1% NP40, protease inhibitor cocktail (Roche, #11836153001), 1 mM NaF, 1 mM Na_3_VO_3_, 1 mM PMSF, 40 U/mL RiboProtect Hu (Blirt, #RT36)) and incubated 10 min on ice to allow gentle cell lysis and extraction of cytoplasmic complexes, as previously (*10*). All buffers used for cell lysis and washing were prepared using the UltraPure DNase/RNase-free water (ThermoFisher Scientific, #10977035). A total of 1mg of cell lysate protein from combined technical replicates was used in each biological experiment (n=3) for immunoprecipitation of complexes using an antibody specific to TIAR (Bethyl, #A303-613A) or a control antibody of the same isotype (Bethyl, #P120-101) and the Protein G-coated magnetic beads (BIO-RAD, #161-4023), split into several technical replicates. Following overnight incubation at 4°C, the complexes were washed three times in a high salt buffer (50nM Tris-HCl pH 7.4, 1M NaCl, 1mM EDTA, 1% NP-40) and then treated with Proteinase K (Promega, #P8107S) in the proteinase buffer (10mM Tris-HCl pH 7.5, 50mM NaCl, 10mM EDTA) for 20 min at 37°C and 1100rpm. Then, an equal volume of proteinase buffer supplemented with 7M Urea (Merck, #U5378) was added, and incubation was continued for 20min. Next, a 1:1 volume of organic phase from the acidic phenol:chloroform pH 4.5 with isoamyl alcohol 125:24:1 premix (ThermoFisher Scientific, #AM9722) was added, and the mix was transferred to the PhaseShield Gel tubes (Life Science Biotechnology, #M2302-05H) for a 5min incubation at 30°C and 11000rpm. After centrifugation (16000xg, 5min, RT). To extract proteins, the same volume of chloroform was added for 5min, and RNA was precipitated from the aqueous phase in the presence of 3 volumes of cold 99.8% EtOH and GlycoBlue Coprecipitant (ThermoFisher Scientific, #AM9515) overnight at-80 °C. After two washes in ice-cold 70% EtOH, the RNA pellet was air-dried and resuspended in water.

### Reverse transcription and Real-time PCR (RT-qPCR)

RNA isolated from the whole cell lysates or immunoprecipitated complexes was analyzed as described previously (*27*). Briefly, RNA was transcribed to cDNA using random hexamers (ThermoFisher Scientific, #51709) and SuperScript III Reverse transcriptase (ThermoFisher Scientific, #18080-044), and after dilution in H_2_O was subjected to the polymerase chain reaction (PCR) using specific primers (Table S2) and the iTaq Universal SYBR Green supermix (Bio-Rad, #1725121) in two technical replicates, following the manufacturers’ protocol. Changes in fluorescence upon SYBR Green intercalation to DNA were monitored using the CFX Connect Real-Time PCR Detection System from Bio-Rad. The Ct values from replicates were averaged, normalized to the Universal Spike I RNA internal control (TATAA Biocenter, #RS25SI) added to each RNA sample, and compared to the reference variant (untreated or isotype IgG). The change in RNA level was quantified using the ΔΔCT method.

### Animals

Mice NOD.Cg-Prkdcscid Il2rgtm1Wjl/SzJ (NSG) 8-10 weeks were from the Jackson Laboratory (JAX#005557). Animals were housed in a temperature-and light-controlled animal facility under a 12h light/12h dark cycle and were provided with standard food and water *ad libitum*. Mice were kept in individually ventilated cages, and all handling procedures were performed under a laminar flow chamber while wearing a sterile surgical disposable gown, cap, shoe cover, and gloves. The mice were treated with a sterile solution of IM (100 mg in physiological salt with 30% PEG-400/kg) or vehicle (30% PEG-400 in physiological salt), daily injected intraperitoneally. The First Local Ethics Committee for Animal Experimentation in Warsaw approved the experimental procedures involving mice, in accordance with EU regulations.

### Ossicles

Formation of ossicles followed a procedure described elsewhere (*42*), with some modifications. To this end, 2×10^6^ of MSC suspended in the mixture of Human Platelet Lysate (Merck, #SCM151) and Matrigel (Merck, #ECM625) were injected subcutaneously in a final volume of 200 μL. After 3 days, the human parathyroid hormone (1–34) (Tocris, #3011/1) daily treatment (40 μg/kg) was introduced for 28 consecutive days. Next, 8 weeks after the injection of hMSC, busulfan (#B2635) 30 mg/kg was administered around 48h before the injection of 2×10^6^ CML cells in physiological salt solution. After 12 weeks, treatment was started with IM for 14 consecutive days. Eventually, tissue samples were collected: blood was retrieved from the heart; spleen and liver were homogenized using a glass douncer and pressed through a 40 μm cell strainer (BD #352340); bone marrow was harvested by flushing from the femur and tibia bones. After washing in PBS, the ACK lysis buffer was added (ThermoFisher Scientific #A1049201) for 10 min at RT, followed by washing in PBS. Then the surface proteins were stained and analyzed by flow cytometry.

### Scaffolds

Cylindrical scaffolds were used to provide spatial support for osteoblast-differentiated cells. The composite films were prepared using a solvent casting method as previously described (*67*) (*68*). Briefly, poly(ε-caprolactone) (PCL; Sigma-Aldrich) was combined with tricalcium phosphate (TCP; particle size ≈ 2.5 µm, Progentix) to obtain homogeneous composite formulations. Scaffolds were fabricated using a precise extrusion deposition technique with a Bioscaffolder device (SYSENG, Germany). Cylindrical scaffolds (7 mm in diameter and 4 mm in height) with a central channel (1.9 mm in diameter) were 3D-printed with a 0.33 mm nozzle. Filaments were deposited in a 0°/90° lay-down pattern between adjacent layers, with a shift equal to half of the theoretical pore size (0.325 mm) introduced in every second layer with the same fiber orientation (n+2 layer), as previously reported (*67*), resulting in scaffolds with an average pore size of approximately 650 µm.

Human osteoblast cell line (hFOB 1.19, ATCC, #CRL-3602) was expanded in DMEM/F12 medium without phenol red (ThermoFisher Scientific, #11039021) supplemented with 10 % FBS (EuroClone, #ECS0180LH) and 0.3 mg/ml of geneticin sulphate (G418, Gibco, UK) at 34 °C. The hMSC were expanded in Minimum Essential Medium alpha (MEM α; ThermoFisher Scientific, #11811023) supplemented with 10% FBS and 1 ng/ml human basic fibroblast growth factor 2 (Merck, #F0291) at 37 °C. The cells were combined at an hFOB:hMSC ratio of 6:4 and then seeded onto the sterilized scaffolds using a droplet containing 5 x 10^5^ of cells. Following 2 h at 37 °C incubation, a mixture of osteogenic media was added composed of one volume of hMSC osteogenic medium (MEM α, 10 mM β-glycerophosphate (β-GP; Merck, #G9422)), and one volume of hFOB osteogenic medium (DMEM/F12, 2 mM β-GP), and supplemented with 10% FBS, 50 μg/ml L-ascorbic acid 2-phosphate (Merck, #A8960), 10 nM of vitamin 1α,25-dihydroxy-vitamin D3 (Merck, #D1530), and 10 nM of dexamethasone (Merck, #D4902). The medium was changed every 2-3 days, and the cells were cultured for 28 days at 37 °C. On the day of scaffold implantation in the mouse, 2×10^5^ K562/luc were added to the center of the scaffold, suspended in 20 μl of Human Platelet Lysate and Matrigel. After 3 days, IM or vehicle treatment was initiated and continued for 14 days.

### Xenotransplants

A mixture of Matrigel with Human Platelet Lysate containing 2×10^6^ of MSC and with or without 0.2×10^6^ K562/luc cells was injected subcutaneously in a final volume of 200 μl. Treatment with IM or vehicle was initiated after 7 days and continued for 14 consecutive days.

### Leukemia cells visualization *in vivo*

To detect K562/luc cells in a mouse, D-Luciferin (Synchem, #103404-75-7) in PBS without Ca^2+^ and Mg^2+^ solution (150mg/kg) was injected 8 min before imaging of 2% isoflurane anesthetized mice in the Bruker In-Vivo Xtreme apparatus. The bioluminescence signal was acquired for 30 sec before the X-ray image of the mouse was taken.

### Tissue staining

Isolated ossicles, scaffolds, or xenotransplants were fixed in 4% buffered paraformaldehyde. The specimens dehydrated in 70%, 96% and 99,8% Ethanol solutions (POCH) were cleared in xylene (Xylene/Mixture of isomers/98,0%, POCH Basic, #BA0860119). After cryoprotecting in 30% sucrose (BioShop, #SUC507.1), the tissue was embedded in the optimal cutting temperature (OCT) compound (Leica, #14020108926) for cryo-sectioning into specimens of 10 μm thickness using a Cryostat Microm HM550 and placed on the microscope slides (ThermoFisher Scientific #J800AMNZ). For brightfield microscopy, immunohistochemistry staining was performed using ImmPRESS Duet Double Staining Polymer Kit (Vector Laboratories, #MP-7724-15) with secondary antibodies anti-mouse IgG conjugated to horse radish peroxidase (HRP) and/or anti-rabbit IgG conjugated to alkaline phosphatase (ALP). Before antibodies, the horse serum (ThermoFisher Scientific #26050-088) was used for blocking, followed by a 2h incubation at RT with antibodies specific to human CD45 (Abcam #ab10559) and human CD31 (Abcam #ab9498). Hematoxylin was used to label the cell nucleus (Shandon, Harris Hematoxylin Non-Acidified, 6765001), and the specimens were clarified in Xylen (POCH) and mounted with medium (Shandon, Consul-Mount Histology Formulation, 9990440). For fluorescent microscopy, the normal goat serum (Abcam #ab7481) was used, and secondary antibodies were: goat anti-rabbit conjugated to FITC and goat anti-mouse conjugated to AlexaFluor-555. Then, cell DNA was labeled with DAPI present in the Vectashield antifade mounting medium (Vector Laboratories, #H1800). Modified Mayer’s Hematoxylin (ThermoScientific, #72804), Eosin-Y Alcoholic (ThermoScientific, #6766007), Alizarin red S (Merck, #1062780025), Oil Red O (Merck, #O0625) were carried out at the Nencki Institute Histology Unit in the Laboratory of Electron Microscopy. Images were collected using the Nikon Eclipse Ti with 40x/0.6 or 20x/0.45 Nikon lenses. Ossicle specimens stained for CD45 detected with ALP were tile-scanned using the Leica DMI 8 microscope and 10x objective. Two sections of each ossicle (n=5) were imaged, and signal intensity was measured within the area of the ossicle using FiJi software.

### Tissue dissociation and flow cytometry cell sorting

The resected tissue fragments were transported in the Hank’s Balanced Salt Solution with calcium and magnesium (ThermoFisher Scientific #14065049), and then were mechanically crushed (cut into pieces with a sterile scalpel), suspended in DMEM/F12 medium heated to 37°C and supplemented with enzymes from the Tumor Dissociation Kit (Miltenyi Biotec # 130-095-929) according to the manufacturer’s instructions, incubated at 37°C for 1h with mixing set to 150 rpm, and pressed through a 0.40 μm cell strainer (BD #352340). Then the ACK erythrocyte lysis buffer was added for 1 min incubation at RT. Following the washing in cold Hank’s medium and PBS, the cells were labeled for the surface protein CD45 using a specific antibody conjugated to BV421, and dead cells were stained using the eBioscience Fixable viability dye (FVD) with eFluor780 (ThermoFisher Scientific, #65-0865-14), following previously described protocols. The K562/luc cells that were FVD-eFluor780 negative, but GFP and CD45-BV421 positive, were sorted using the BD FACS Aria (RRID:SCR_018934) and used for RNA isolation or determination of CD proteins.

### RNA-seq and data analysis

RNA was isolated from K562 cells, collecting suspension cells from a co-culture with HS-5 that was initiated under hypoxia 48h before treatment with 1 μM IM for 18h; or from K562/luc cells sorted by FACS from xenotransplants. RNA was isolated from whole cells, and DNA was removed by and on-column DNase treatment as previously described in detail (*27*). Samples were qualified for sequencing if the RNA integrity number was larger than 8.5. During library preparation, poly(A+) RNA was selected and prepared for RNA-seq as described previously (*27*). Samples were sequenced using the paired-end 150–base pair (bp) reads on Illumina NovaSeq 6000 instrument (RRID:SCR_016387) at Yale Center for Genome Analysis core facility. Preprocessing of datasets, analysis of differential gene expression, intron splicing, and alternative splicing were performed as previously (*27*). Briefly, adapters were trimmed using FASTP, and reads were mapped to the hg38 genome using the STAR aligner (v2.7.7a). Then the featureCounts program (Subread/2.0.3) output was used to count the number of reads mapping to each gene (statistics presented in Table S3), followed by differential gene expression analysis using the R/Bioconductor package DESeq2 (v1.40.2) in RStudio (v4.3.1), and applying a variance stabilizing transformation (vst function) with the blind argument set to FALSE. Significantly different genes with log2 fold change of expression between two variants ≥ 0.6 were considered upregulated, and with ≤ 0.6 downregulated. The Python tool SPLICE-q was used for the quantification of intron splicing (SPI), and rMATS-turbo to analyze alternative splicing, with the same settings described before (*27*). Comparison of the functional annotation enrichment analysis of expressed genes (at least 10 reads mapped in 3 replicates and ranked by Wald statistic from DESeq2) was performed using the R package Gene Set Enrichment Analysis (GSEA) (*69, 70*), with minSize = 15 and maxSize = 500, and the Molecular Signatures Database (MSigDB) v2026.1.Hs as a reference. The MSigDB C2 collection (Curated gene sets) was used for Reactome, and C5 (ontology gene sets) for Gene Ontology (GO) Biological Processes (BP) and Molecular Functions (MF) enrichment analysis.

### Analysis of published RNA-seq data

Data from rMATS-turbo analysis of RNA alternative splicing in samples of CD34+ enriched cells obtained from bone marrow biopsies collected before and after the patient’s 6-month treatment with IM are available through the National Center for Biotechnology Information (NCBI) under GEO accession number GSE310243 (combined samples SRR36072325 and SRR36072321 compared to combined samples SRR36072320 and SRR36072324) and described before (*27*). The gene expression changes analyzed by single-cell RNA sequencing (scRNA-seq) in bone marrow aspirates obtained from healthy donors and CML patients were from the Single-cell atlas of diagnostic Chronic Myeloid Leukemia bone marrow (scdbm) (http://scdbm.ddnetbio.com) (*45*). Only results for CD34+ cell subtype are presented for each of the categories that included samples from patients classified into the following categories: Control (healthy donors; n=8), A (CML patients with good response to imatinib; n=9), B (CML patients with IM failure; n=9), and C (CML patients with pan-TKI resistance and progression to blast crisis stage; n=9).

### Flow cytometry analysis

Apoptosis level (AnxA5), cell cycle progression, CD surface proteins, and metabolic features stained following the dedicated protocols were analyzed using BD LSR Fortessa Analyzer (RRID:SCR_018655) equipped with the 355 nm, 405 nm, 488 nm, and 635 nm lasers. FlowJo version 10.10 software (RRID:SCR_008520) was used to analyze the flow cytometry results.

### Fixation viability dye (FVD), AnxA5 and cell cycle

Dead cells were stained before cell fixing, by incubating the cells in cold PBS for 20 min with the Fixable Viability Dye from ThermoFisher Scientific, selected depending on the fluorescent properties: eFluor455UV (#65-0868-14), eFluor660 (#65-0864-14), or eFluor780 (#65-0865-14). Apoptosis level was estimated with AnxA5-FITC (BD Pharmingen #556419) and 7-AAD (BD Pharmingen #559925) staining in the Annexin V Binding Buffer (BD Pharmingen #556454), and cell cycle progression was monitored based on cell DNA content stained with PI (BD Pharmingen #556463) and analyzed with ModFit software following previously described protocols (*10*).

### Cell differentiation (CD) surface markers

To stain surface markers 3×10^5^ cells were washed in cold PBS and resuspended in the Brilliant Stain Buffer (BD Horizon #563794). Following a 10 min incubation at room temperature with human Fc Receptor Binding Inhibitor (Invitrogen #14-9161-73), the cell surface markers were stained at 4°C for 30 min with antibodies conjugated to fluorochromes: CD235a-BB515 (BD Horizon #565233) or CD235a-BV605 for GFP-positive cells (BD Horizon #740409), IgG2b isotype control to CD235a-BB515 (BD Horizon #564510), CD45-BV-421 (BD Horizon #563879), CD33-PerCP (BD #333146), CD36-APC (Biolegend #336207). After two washes in PBS with 1% bovine serum albumin (Sigma-Aldrich #A3294), the cell pellet was resuspended in PBS and added to equal volume of 4% paraformaldehyde for 15min incubation at room temperature before cell analysis.

### Mitochondrial Membrane Potential

The cells were seeded in co-culture hypoxic conditions 48h before 1 μM IM was added for 24h. For the assay, 2×10^5^ of K562 cells were suspended in 100 μl of warm complete growth medium with or without 10 μM CCCP (Sigma-Aldrich #C2759) for 5 min at RT (handling in hypoxia workstation). Then an equal volume of warm growth medium (placed under hypoxia in advance to adjust oxygenation and heated to 37°C) with 3 μM JC-1 probe (Thermo Fisher Scientific, #T3168) was added (and w/o 10 μM CCCP), and the cells were incubated for 25min at 37°C under hypoxic conditions. After one washing in warm PBS the cells were suspended in PBS. Fluorescent signal of JC-1 probe was detected in two channels: red dimers in mitochondria – in BV605 (with band pass 605/12nm) and green monomers in cytoplasm – in AlexaFluor 488 (with 530/30nm filter).

### ROS level

The 2’,7’-dichlorodihydrofluorescein diacetate (H_2_DCFDA) cell-permeant probe was used to determine the cellular level of reactive oxygen species (ROS). The cells were seeded 48h in co-culture hypoxic conditions before adding IM at the final concentration of 1 μM for 24h. As a positive control, 100 μM H_2_O_2_ for 1h was added. Then 10^6^ of K562 cells were washed in warm PBS (placed under hypoxia in advance to adjust oxygenation and heated to 37°C) and incubated in PBS with 2 μM H_2_DCFDA (ThermoFisher Scientific, #D399) for 30min at 37°C under hypoxia. After 2 washes in warm PBS, the cells were stained with 7-AAD (BD Pharmingen #559925) for 2 min before the measurement. From each sample, 10^3^ of the 7-AAD-negative cells were recorded.

### Reduction potential

The cells were seeded in co-culture hypoxic conditions 48h before 1 μM IM was added for 24h. Reduction potential of cells was determined using the RealTime-Glo MT Cell Viability kit (Promega, #G9711) following the manufacturer’s protocol. Then, the same number of live K562 cells (4×10^3^) was used for the experiment, and the volume of cell culture was equalized to 20 μl between the samples using warm growth medium (placed under hypoxia in advance to adjust oxygenation), placing the tubes in a warm holder. Then 20 μl of warm growth medium (equal volume) with enzyme and a substrate diluted 1000x (premix for all samples) was added for 30 min of incubation at 37 °C. The luminescence signal was detected using GLOMAX Luminometer (Promega) with integration time set to 1 sec.

### Oxidative DNA and RNA damage

The cells were seeded in co-culture hypoxic conditions 48h before 100 μM H_2_O_2_ was added for 1h. First, 5×10^5^ of K562 cells were washed in cold PBS before fixation in 2% paraformaldehyde (MP Biomedicals #30525-89-4) for 15min on ice. After washing in PBS, 10min of permeabilization on ice in 0.05% saponin (Calbiochem #558255) in PBS, washing in PBS, 30 min of blocking on ice in 3% normal goat serum (Abcam #ab7481), the cells were centrifuged and resuspended in the residual PBS (about 50 μl). Then 50 μl of staining buffer was added (0.05% saponin in PBS) with 1 μl of 8-OXO-FITC antibody (Abcam #ab183393) for 1h incubation on ice. After two washes in 0.05% saponin in PBS, the cell pellet was resuspended in PBS for flow cytometry analysis.

### Lipid peroxidation

The cells were seeded in co-culture hypoxic conditions 48h before 1 μM IM was added for 24h. The lipid peroxidation (LOOH) was checked using the Click-iT Lipid Peroxidation Detection with Linoleamide Alkyne (LAA) kit (ThermoFisher Scientific, #C10446) following the manufacturer’s protocol. First, 1×10^6^ of K562 cells were suspended in growth medium (placed under hypoxia in advance to adjust oxygenation and heat to 37°C) supplemented with LAA at a final concentration of 50 μM for a 4h incubation at 37°C under hypoxia. After washing in PBS, the cells were fixed in 2% PFA in cold PBS and incubated 15 min at RT, then washed in PBS and permeabilized in 0.25% Tween-20 (SigmaAldrich, #P9416) in PBS for 15 min at RT. Following blocking in 3% BSA and washing in PBS, the cells were incubated for 30 min at RT in the Click-it reaction cocktail containing CuSO_4_ and Azide-488 and then washed in PBS before flow cytometry analysis.

## Statistical analysis of results

All procedures were executed for three independent biological experiments, unless stated otherwise in the Figure legend. Data were presented as the mean ± SD or range. Statistical analysis was performed using GraphPad Prism version 9.3.1 (RRID:SCR_002798). The unpaired two-way Student’s t-test was used to compare two samples with the same variation, and p<0.05 was considered statistically significant and designated by: *p≤0.05; **p≤0.01; ***p≤0.001; ****p≤0.0001. The nonparametric two-tailed Mann-Whitney test was used to compare results from *in vivo* experiments, and the exact significance values are presented in the plots. Two-way ANOVA was used to compare the selected variant (@) with several other samples.

Graphics in the schematic presentations of the experimental workflow were created at BioRender.com.

## AVAILABILITY OF DATA AND MATERIALS

Realization of the project did not involve the generation of new materials. The proteomic data from this publication have been deposited to the ProteomeXchange Consortium via the PRIDE partner repository with the identifier PXD032332. Raw RNA-sequencing data are available through the National Center for Biotechnology Information (NCBI) under GEO accession numbers GSE332859.

## ACKNOWLEDGEMENTS

We would like to thank members of the Cytometry Laboratory at Nencki Institute for assistance with tissue collection and flow cytometry analysis, in particular Julian Swatler, Agata Kominek, Lukasz Bugajski, and Marta Olchowik. We would like to thank members of the Animal House at the Faculty of Biology for assistance with mouse breeding. We are grateful to members of the Neugebauer laboratory for helpful discussions. This work was supported by a grant from the National Science Center in Poland (2018/30/M/NZ3/00274 to P.P.-B.). We also thank the National Institutes of Health (NIH/NCI 2P30CA006927-56A1 to T.S; R01GM140735 and R01GM112766 to KM.N.) and the National Science Center in Poland (2021/41/B/NZ5/04077 to K.P; 2020/39/I/ST5/03473 to W.S.) for support. P.P.-B. received additional support from a Senior Fulbright Award (2022–23) from the Polish-US Fulbright Commission. P.P.-B. is a Forbeck Scholar (2024-27) and a member of the “TRANSLACORE” Cost Action CA21154. Data acquisition at Yale Center for Genomic Analysis was supported by the National Institute of General Medical Sciences of the National Institutes of Health under award number 1S10OD030363-01A1. The contents of this article are solely the responsibility of the authors and do not necessarily represent the official views of the NIH.

## AUTHOR CONTRIBUTIONS

P.P-B. and E.K. conceptualized the study; P.P-B., M.W. and BV.L. carried out laboratory experiments with cell culture; P.P.-B. carried out flow cytometry analysis; P.P-B., E.K. and P.P-K. carried out procedures on mice; E.W and J.I. prepared the scaffolds; P.P-B. and L.S. analyzed the RNA-seq data; R.S. performed protein mass-spectrometry and quantitative data analysis; A.K. did cryo-sectioning; A.K., P.P-B. and A.M-P. did histological staining; W.S. supervised J.I. and E.W.; T.S. supervised BV.L.; K.P., T.S., W.S. and KM.N. consulted the results; K.P. and KM.N. mentored P.P-B. The original manuscript was written by P.P-B. and was subsequently edited by R.S., J.I., L.S., M.W., KM.N. and K.P. The project was supervised by P.P.-B.

