## Supplementary Material for "TIAR-dependent coordination of alternative splicing and lipid peroxidation is required for CML cell resistance to imatinib in the bone marrow stroma"

### Supplementary materials

This PDF file includes:

Figures S1-S14 with text legends
Tables S1-S3 with text legends

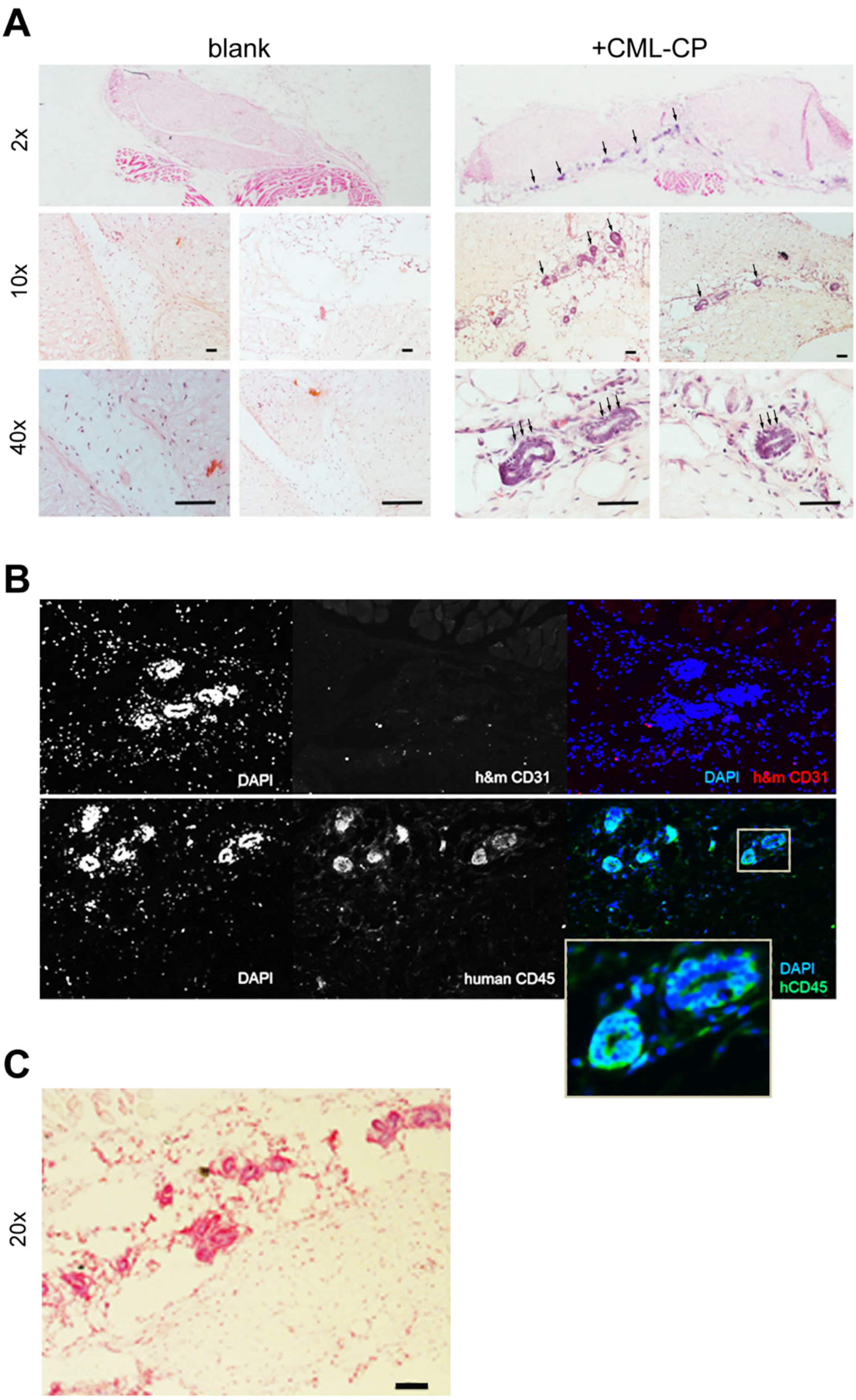

**Figure S1. Detection of human leukemia cells in ossicle specimens.**

**(A)** Eosin and hematoxylin staining of ossicles from mice without leukemia cells injection (blank) or having injected Lin-CD34+ cells from CML patient (+CML-CP). Arrows indicate concentrated cells with stronger nuclear staining detected in ossicles with leukemia cells under a bright light microscope **(A)** and fluorescence microscope with DAPI staining **(B)**. **(B)** Immunostaining of CD45 and CD31 proteins of human (h) or mouse (m) origin. Representative images of fluorescence intensity detected in each channel are presented in gray scale, channels overlay in color, enlarged region is marked with a square. **(C)** Immunostaining of human CD45 with an antibody conjugated to alkaline phosphatase, visible in red; bar = 100 µm.

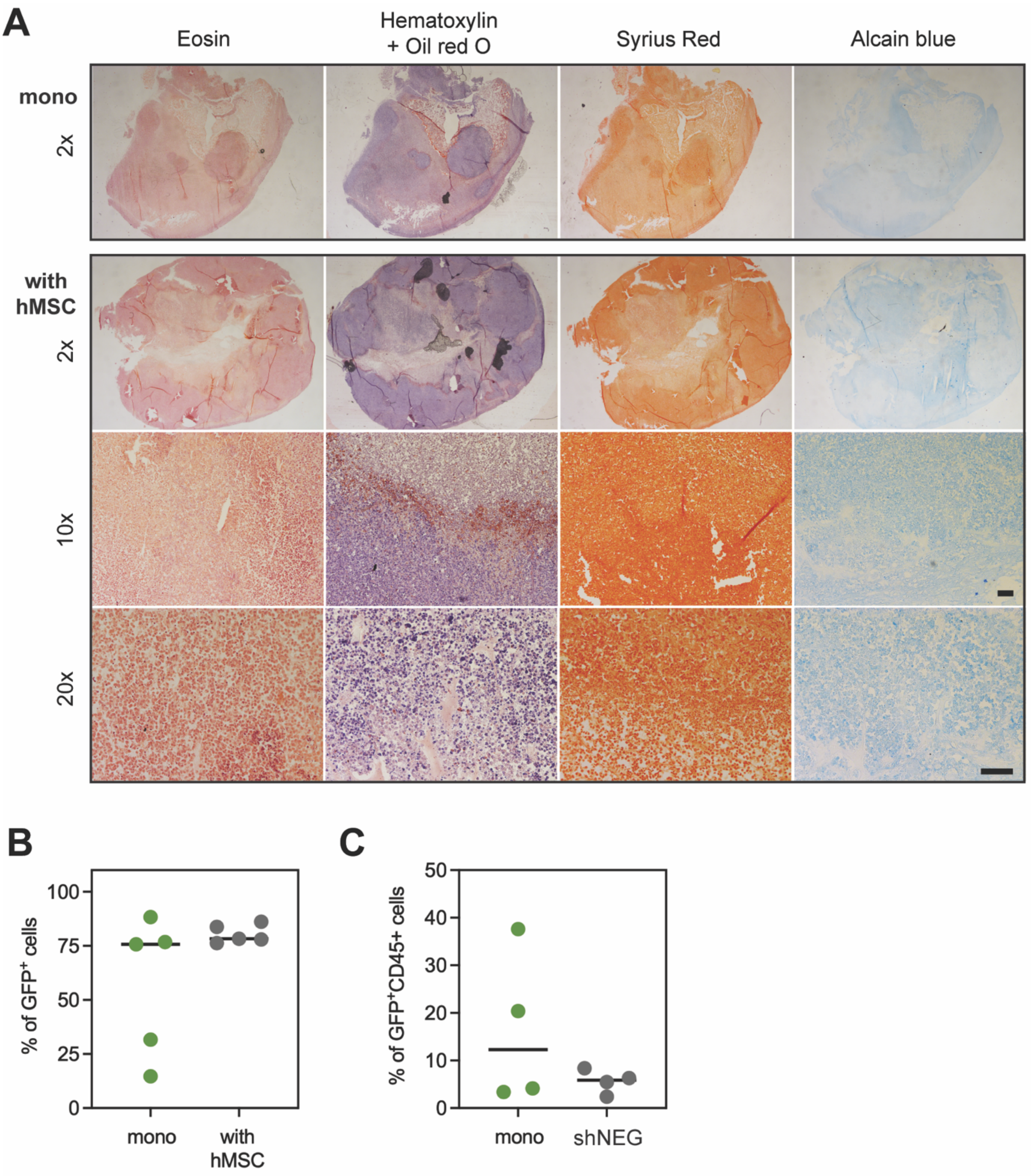

**Figure S2. Characteristics of xenografts formed by K562/luc cells expressing shNEG formed upon injection of leukemia cells mixed with hMSC or without (mono).**

**(A)** Staining of xenograft sections with dyes indicated in the top panel. Imaging with brightfield microscope upright (objective 2x) and inverted (objectives 10x and 20x); bar = 100 µm.
**(B-C)** Percent of live cells (negative for fixable viability dye eFluor780) in xenografts from untreated mice (with hMSC) that express GFP (n=5) **(B)** or treated for the last 14 days with vehicle (shNEG) that are positive for both GFP and human CD45-BV421 staining (n=4) **(C)**, analyzed by flow cytometry.

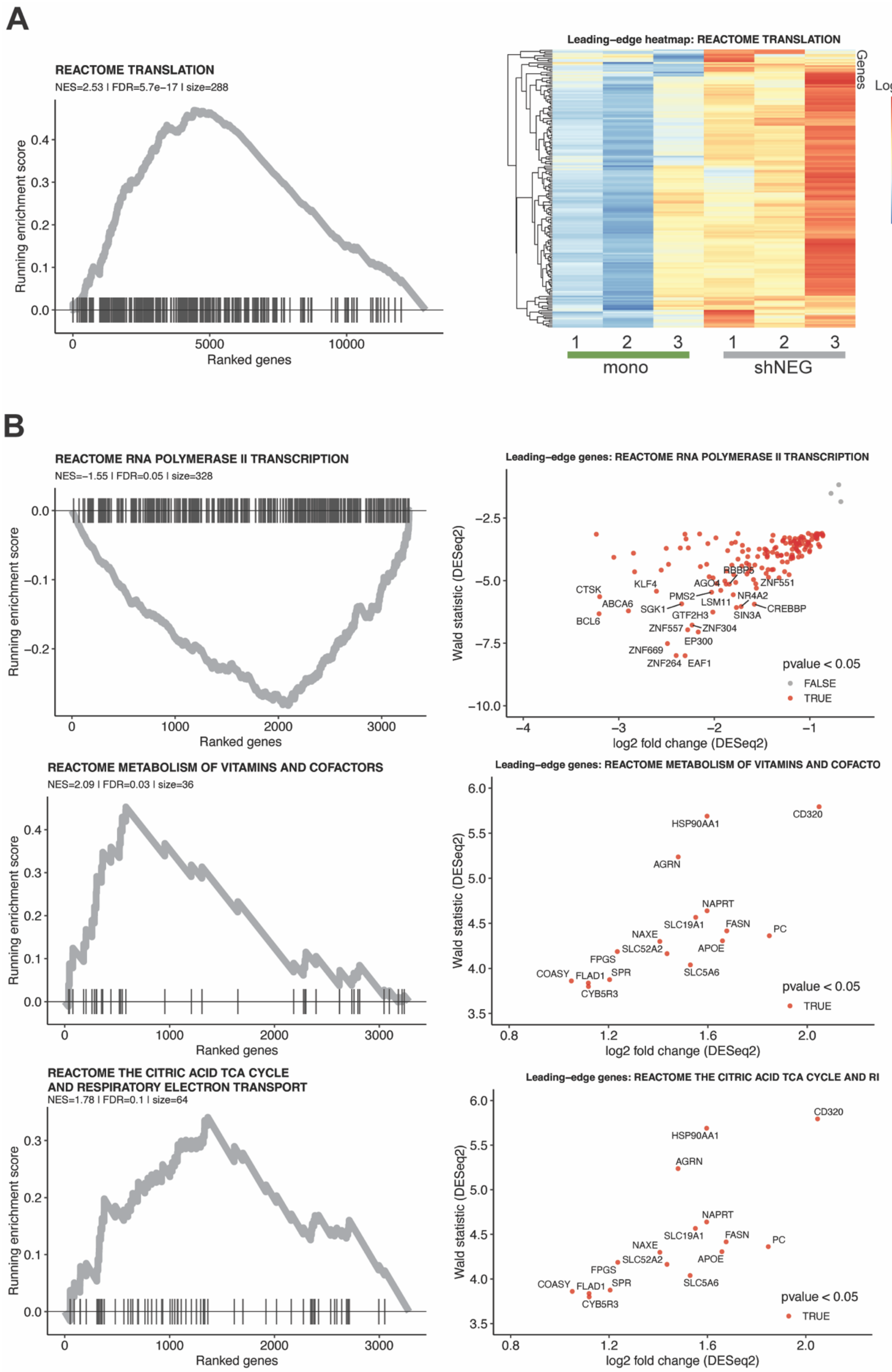

**Figure S3. Enrichment of functional annotation of genes differentially regulated in leukemia cells xenografts with or without human bone marrow stromal cells (legend next page)**

**Figure S3. Enrichment of functional annotation of genes differentially regulated in leukemia cells xenografts with or without human bone marrow stromal cells.**

Analysis of RNA from K562/luc cells (expressing control shNEG, Firefly luciferase, and GFP), FACS-sorted from xenografts formed following injection of the leukemia cells alone (mono) or mixed with human primary bone marrow mesenchymal stem cells and treated with vehicle for the last 14 days (shNEG).
The top Reactome terms identified by the Gene Set Enrichment Analysis (GSEA) of the DeSeq2 results, with the highest normalized enrichment score (NES) and the lowest $P_{adj.}$ (FDR); size - number of genes in the sample annotated per term > 30 required. GSEA analysis of **(A)** the full list of identified transcripts and **(B)** the list of transcripts upregulated (log2 value of fold change ≥ 0.6) or downregulated (log2 value of fold change ≤ -0.6).
**(A-B) Left panel** - The running enrichment score presented by the grey line for each of the ranked genes marked with a vertical black line.
**Right panel** – leading-edge gene expression level as log2 of fold change in xenografts (1, 2, and 3) shNEG versus mono; names of top 15 changed displayed.

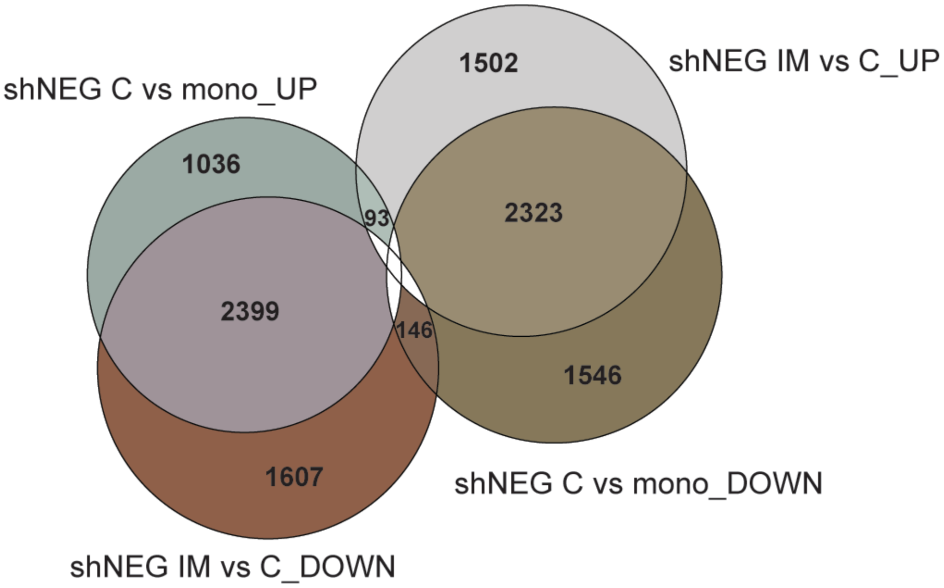

**Figure S4. Comparison of genes expressed in leukemia cells from xenografts.**

Genes expressed in K562/luc shNEG cells FACS-sorted from xenografts formed without hMSC (mono) and with hMSC treated for two weeks with vehicle (shNEG_C) or imatinib (shNEG_IM). Indicated is the number of genes that are upregulated (UP, log2 value of fold change $\geq 0.6$) or downregulated (DOWN, log2 value of fold change $\leq -0.6$) in differential gene expression analysis by DeSeq2.

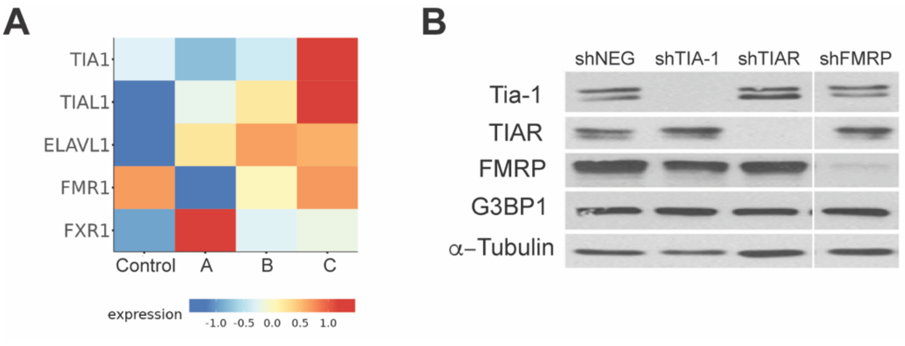

**Figure S5. Expression of selected RNA-binding proteins .**

**A)** Analysis of data at the Single-cell atlas of diagnostic Chronic Myeloid Leukemia bone marrow (scdbm) for CD34+ cells subtype (*43*). Data and detailed description of patient classification available: http://scdbm.ddnetbio.com; A – responded to IM within 12 months; B – IM treatment failed within 18 months; C – resistant to IM and other TKI-s. **B)** Protein level detected by Western blot in whole cell lysates of K562 transduced with shRNA.

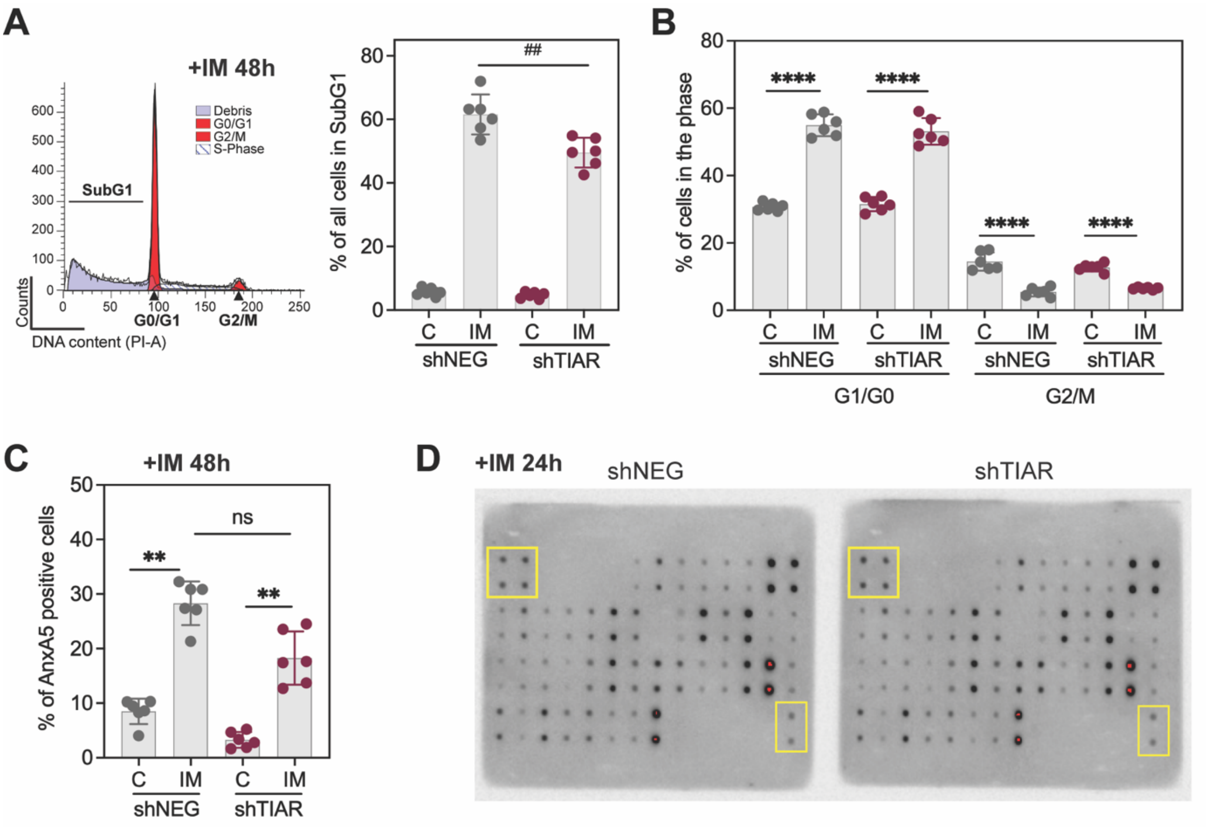

**Figure S6. Influence of TIAR knock-down on the cell cycle progression and cell death upon imatinib treatment.**

Results obtained from the K562 cells expressing shNEG or shTIAR co-cultured with HS-5 cells in hypoxia (1.5% O2) for 72h that were treated (IM) or not (C) with imatinib for 48h **(A-C)** or 24h **(D)** before cell collection for analysis.

**(A-B)** Cell cycle progression analysis based on DNA content in the cells stained with propidium iodide (PI).

**(A)** Right panel – representative image of cell cycle phase identified by fluorescence intensity analysis in ModFit. Left panel – percent of dead cells with fragmented DNA (SubG1).

**(B)** Percent of cells in G0/G1 and G2/M phases.

**(C)** Percent of AnxA5-FITC positive cells.

**(B-C)** Mean value from independent experiments (n=3) ± SD for two different clones of cells expressing each shRNA. Student's two-way t-test was used to compare IM vs C (*) and shTIAR vs shNEG (#); ** $p \leq 0.01$, **** $p \leq 0.001$, ns – $p > 0.05$.

**(D)** Apoptotic signaling antibody arrays detecting proteins in cell lysates; reference dots marked in yellow square; imagining showing signal saturation (in red).

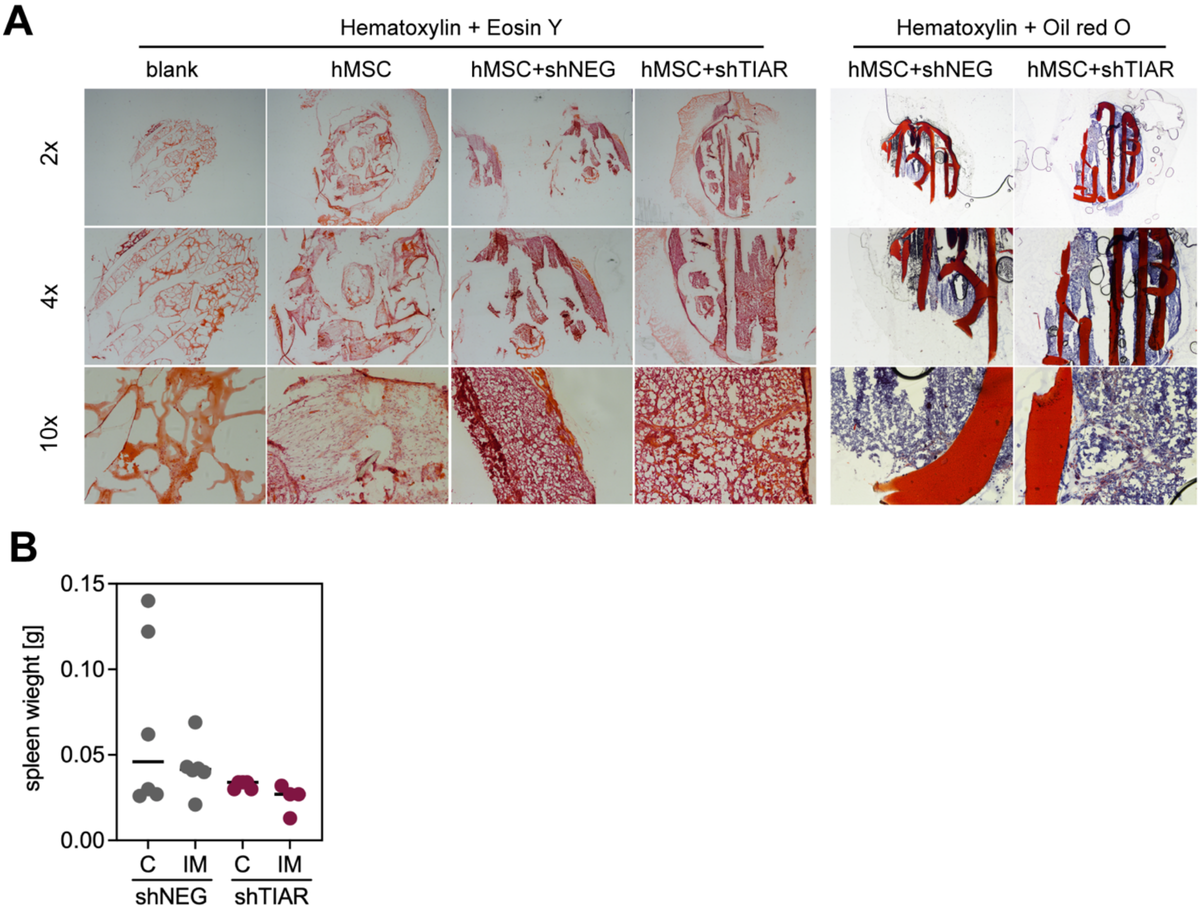

**Figure S7. Analysis of tissue sections and spleen size.**

**(A)** Histological analysis of sections of scaffolds that were seeded before implantation only with hMSC, with a mix of hMSC and K562/luc cells with shNEG or shTIAR, or blank - not seeded with cells before implantation. Staining with dyes is indicated in the top panel. Imaging with a brightfield microscope, upright objective 2x, 4x, and 10x.

**(B)** Weight of spleens isolated from mice (n=dot) with xenografts.

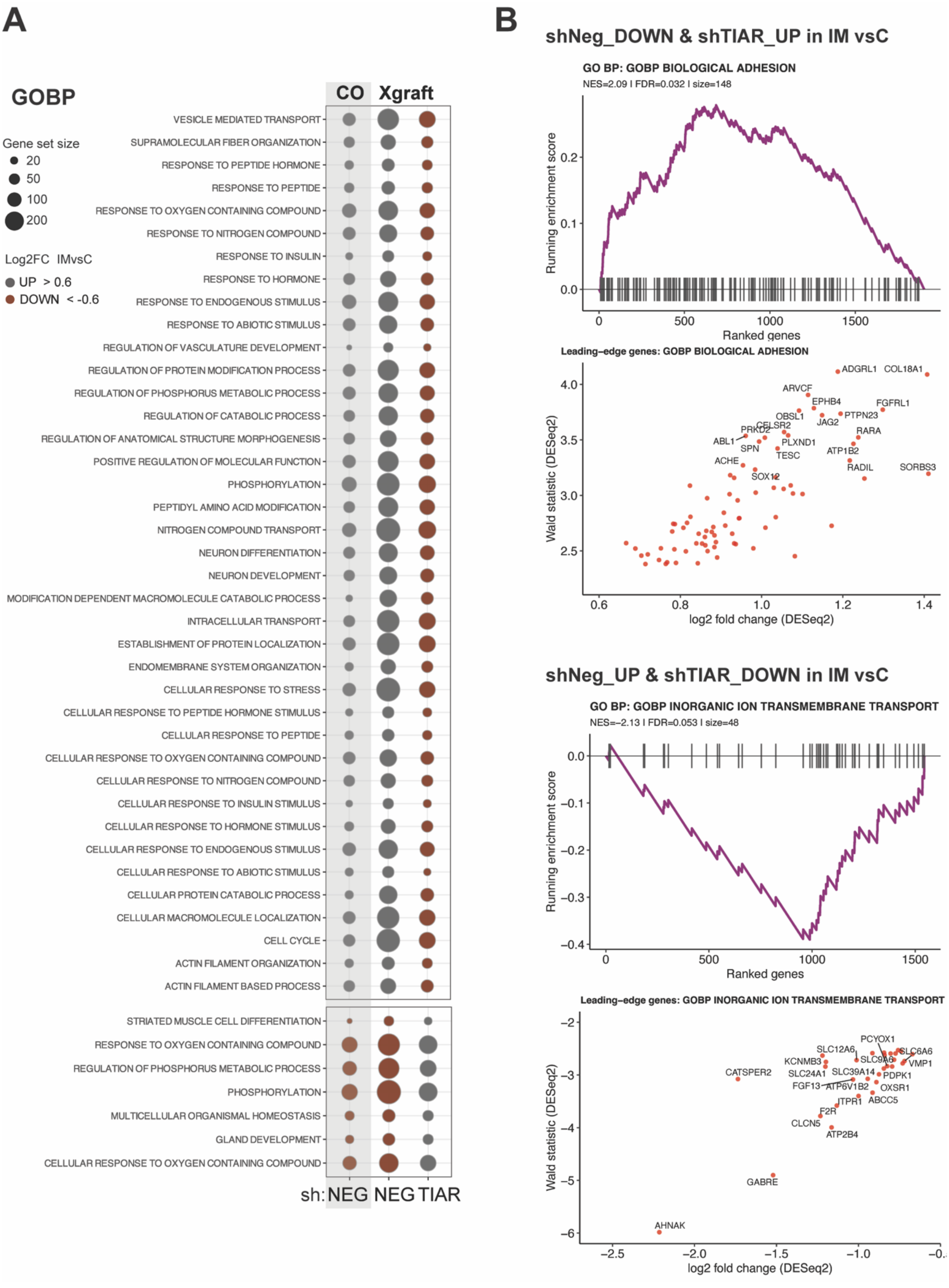

**Figure S8. Functional annotation of genes reversely regulated upon imatinib in cells with TIAR silencing. (legend next page)**

**Figure S8. Functional annotation of genes reversely regulated upon imatinib in cells with TIAR silencing.**

**(A)** Gene Ontology Biological Processes (GOBP) terms identified by the GSEA of genes that upon IM vs C in xenografts (Xgraft) of shNEG are reversely regulated in shTIAR (shaded in purple). In grey circles - UP (log2 of fold change, Log2FC > 0.6) n = 1993, brown circles DOWN (Log2FC < -0.6) n = 1756. Selected terms with $NES_{abs} \geq 1.45$, p-value $\leq 0.05$, and the number of genes in the sample annotated per term (size) $\geq 10$; n -number of genes. Results for selected terms compared to GSEA for UP and DOWN genes in shNEG co-cultured (CO) with HS-5 cells for 48h in hypoxia 1.5% $O_2$ before IM 18h treatment versus untreated.

**(B)** The top GOBP terms identified by the GSEA **(A)**, with the highest normalized enrichment score (NES) and the lowest $P_{adj.}$ (FDR); size - number of genes in the sample annotated per term > 30 required. The running enrichment score for each of the ranked genes (marked with a vertical black line) presented by the purple line. The leading-edge gene expression level as Log2FC in xenografts; names of the top 15 changed with $p < 0.05$ are displayed.

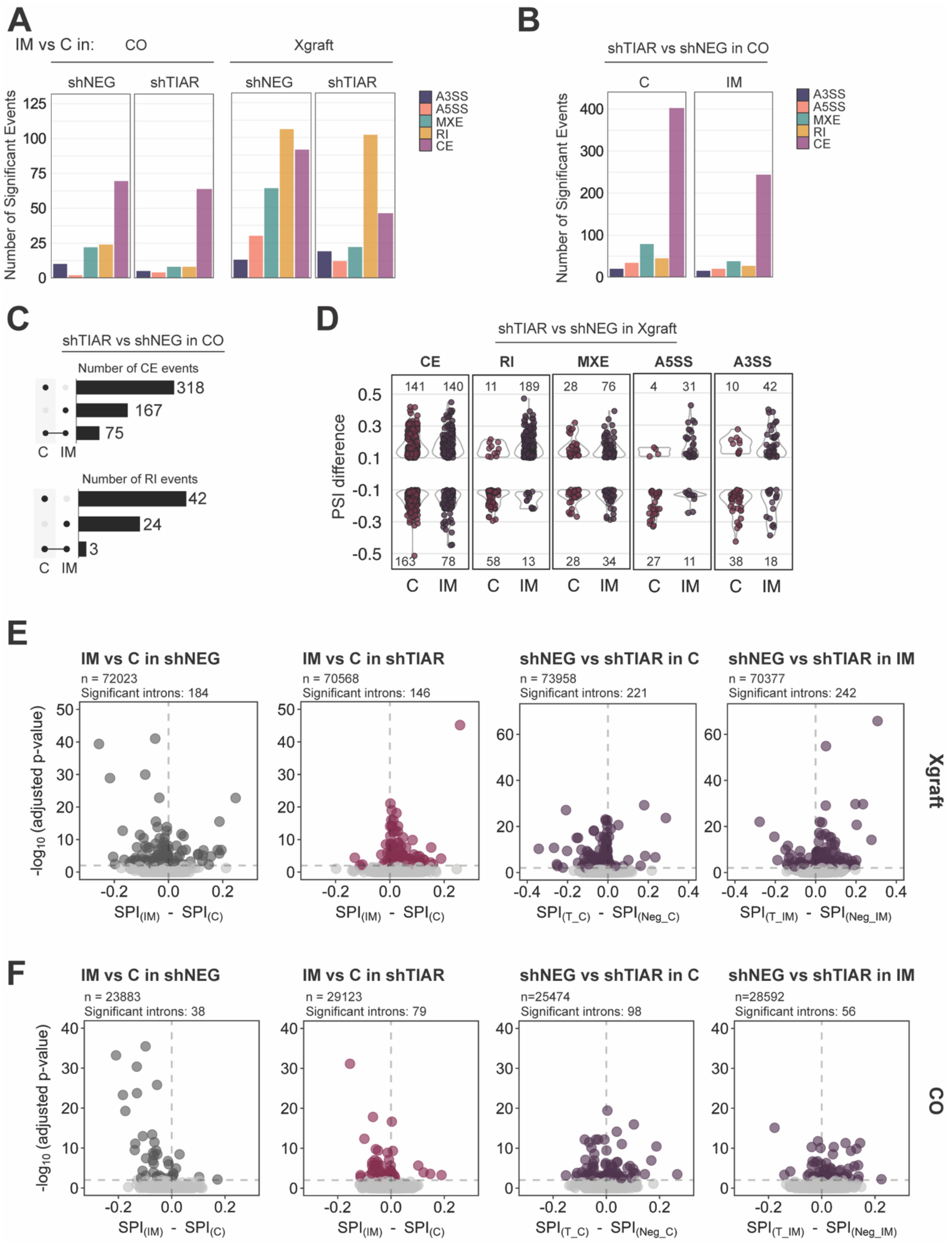

**Figure S9. Changes in splicing upon imatinib treatment and TIAR silencing in different experimental setups. (legend next page)**

**Figure S9. Changes in splicing upon imatinib treatment and TIAR silencing in different experimental setups.**

Presented are results of rMATS analysis of RNA from cells FACS-sorted from xenografts (Xgraft) and co-cultured (CO) that were treated with imatinib (IM) or not (C). Significant events had at least 20 reads, difference in the PSI (Perecent Spliced In) > 0.1 with FDR < 0.05; Alternative splicing (AS) events analyzed: CE – cassette exon, RI – intron retention, MXE – mutually exclusive exons, A5SS – alternative 5' splice site, A3SS – alternative 3' splice site.

**(A)** Number of alternative splicing events that significantly differ in IM vs C in shNEG or shTIAR cells from CO or Xgraft.

**(B)** Number of alternative splicing events that significantly differ in shTIAR vs shNEG cells from CO in IM or C conditions.

**(C)** Comparison of CE (upper panel) and RI (lower panel) splicing events that significantly differ in shTIAR vs shNEG cells from CO in IM or C conditions.

**(D)** PSI difference of AS events that significantly differ in shTIAR vs shNEG cells from Xgraft in IM or C conditions; number of events with PSI >0.1 and <-0.1 provided in the plot.

**(E-F)** Change in SPI values (ΔSPI) in cells isolated from Xgrafts **(E)** or CO **(F)**. Introns (n) detected by at least 500 reads and with significant changes in SPI (adjusted P value < 0.05) are marked in colors. Adjusted P values were determined using chi-square tests for pairs of replicates and the Benjamini-Hochberg procedure.

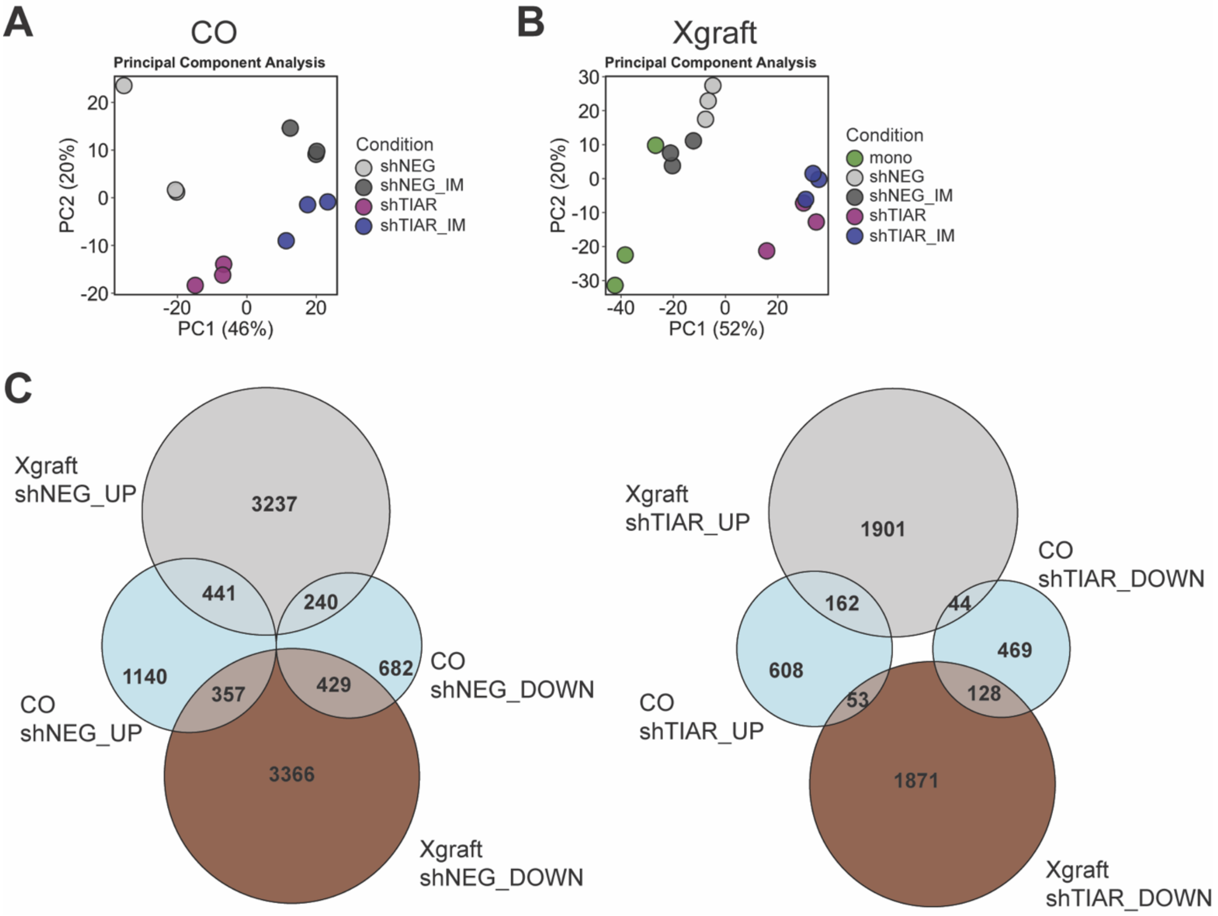

**Figure S10. Genes with expression altered upon imatinib and TIAR knockdown in K562 cells from co-culture and xenografts.**

Results of differential expression analysis of the variant with hMSC versus mono by DeSeq2 of RNA sequenced from FACS-sorted cells. Presented are the results from cells isolated from three different xenografts for each variant (Xgraft) and from a hypoxic co-culture (CO) treated or not with imatinib (IM).

**(A-B)** Principal component analysis.

**(C)** Comparison of genes with expression altered upon IM, so are regulated UP (log2 of fold change, Log2FC > 0.6) and DOWN (Log2FC < -0.6).

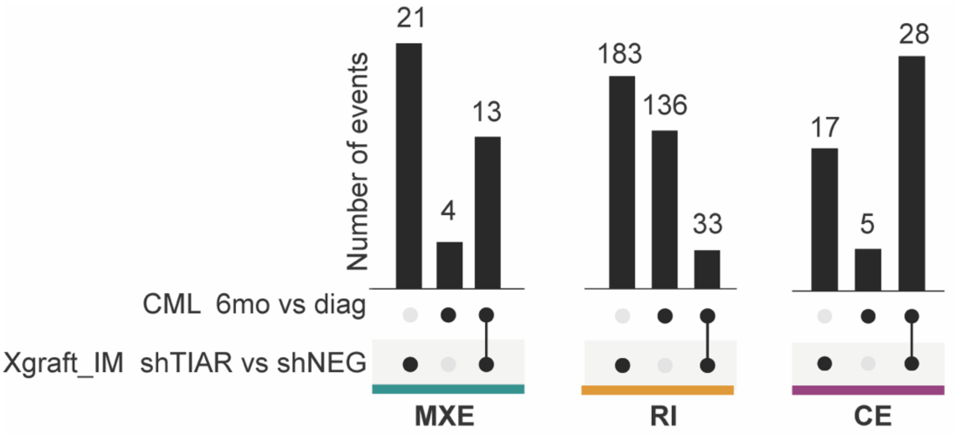

**Figure S11. Alternative splicing events significantly changed in CML cells upon long-term imatinib treatment.**

Analysis of alternative splicing events in xenografts imatinib-treated shTIAR versus shNEG cells (Xgraft_IM) and in CD34 positive cells from bone marrow biopsies of a patient with CML (GSE310243 (*27*)) obtained after 6 months of imatinib therapy versus obtained at the diagnosis (CML 6mo vs diag). Compared only significant events detected by at least 20 reads, with an absolute value of inclusion difference > 0.1, and with FDR < 0.05; MXE - mutually excluded exons, RI – retained intron, CE – cassette exon.

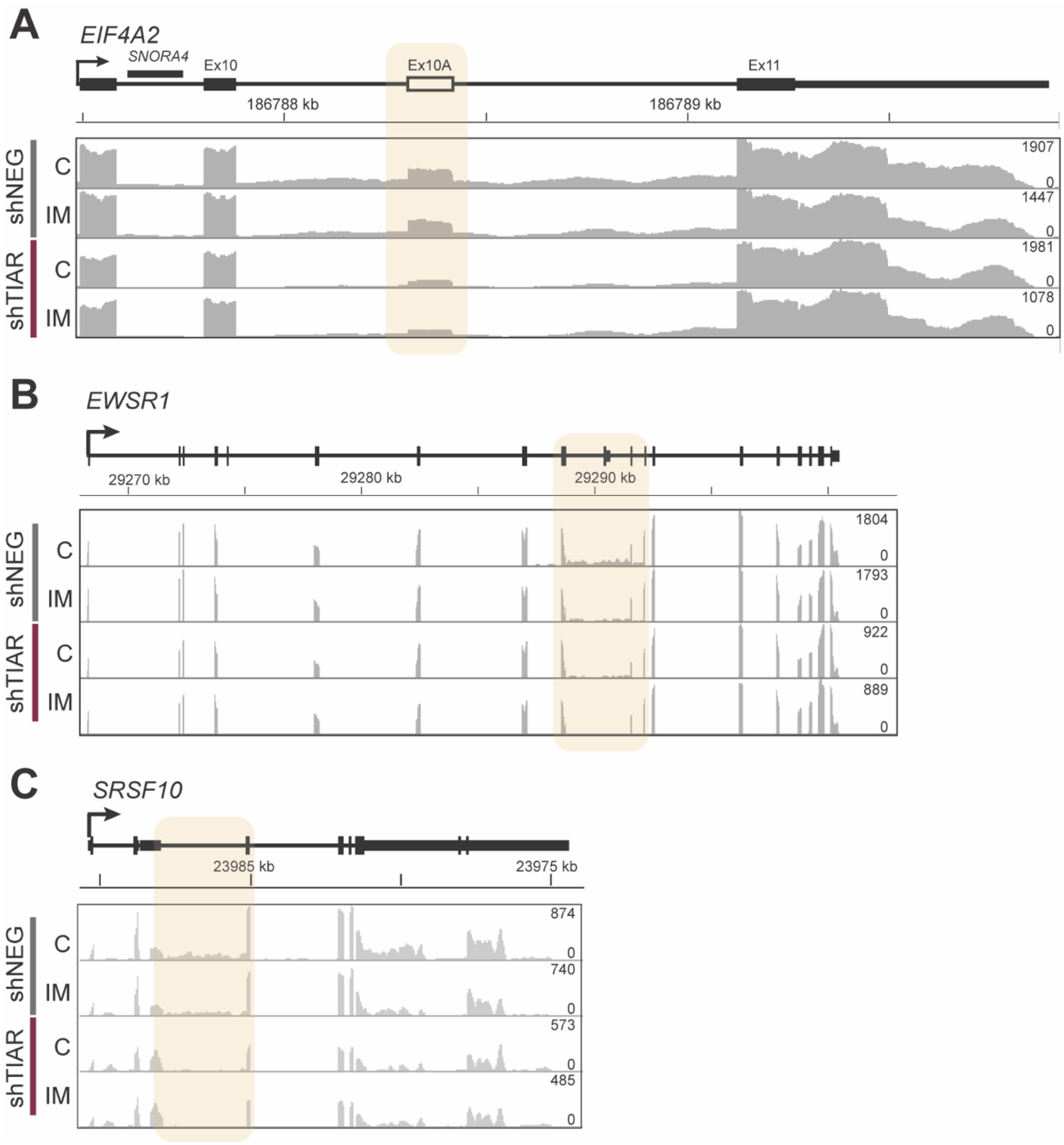

**Figure S12. Examples of genes with altered alternative splicing in cells from xenografts treated with imatinib.**

Gene coverage from RNA sequencing in one of the 3 experiments on K562/luc cells FACS-sorted from xenografts expressing shNEG or shTIAR and treated with imatinib for two weeks (IM) or vehicle (C). Reads alignment to EIF4A2 **(A)**, EWSR1 **(B)**, and SRSF10 **(C)** is presented in grey in the bottom panel, the maximal number of reads in the left corner, and genomic coordinates in the middle panel, a schematic representation of gene regions of exons and introns in the top panel, and yellow shadowing marks regions with changes in alternative splicing.

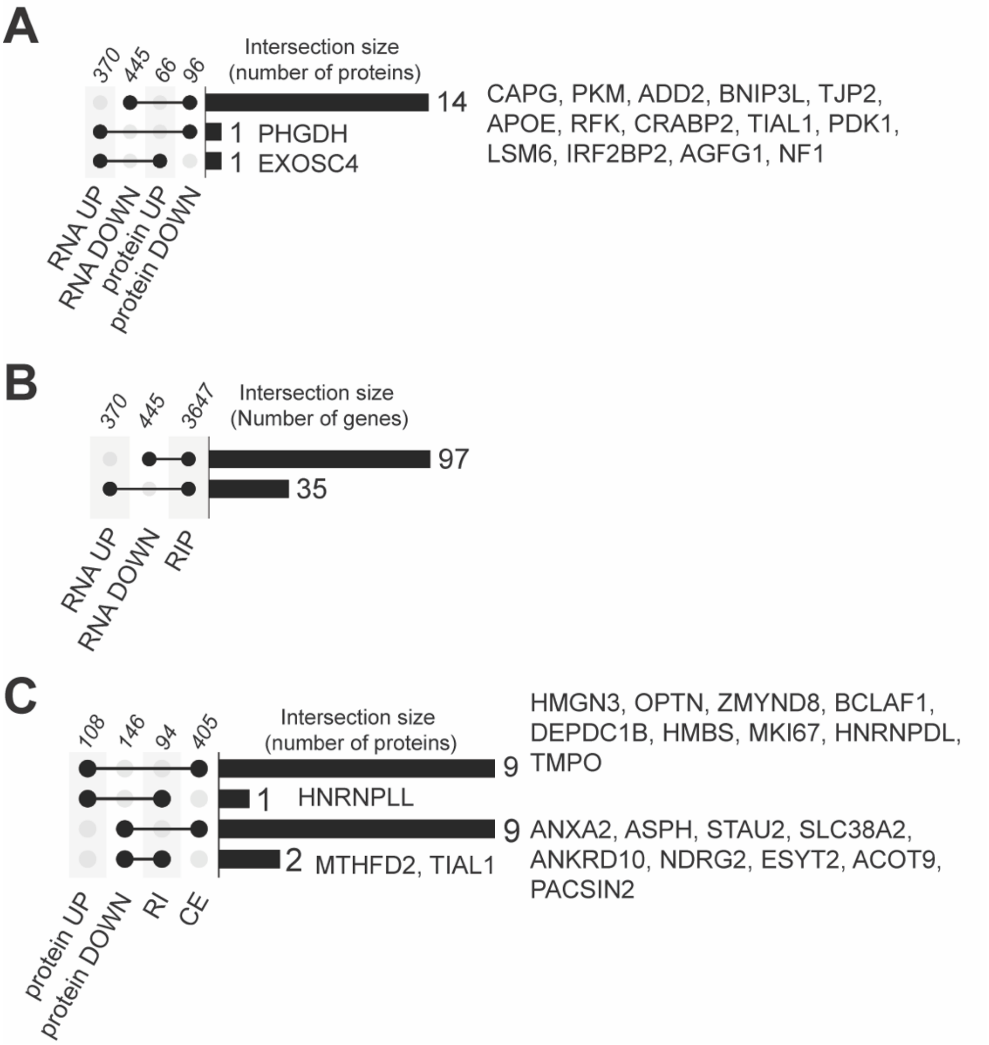

**Figure S13. Comparison of proteins and RNA significantly altered in cells upon imatinib treatment.**

Results of mass spectrometry analysis of nascent protein synthesis (Protein) and short-read sequencing analysis of RNA (RNA) from K562 co-cultured with HS-5 cells in hypoxia 1.5% $O_2$ (CO) for 48h before IM 18h treatment was initiated. Proteins and RNA with shTIAR vs shNEG that had fold change log 2 value ≥ 0.6 (UP), and ≤ -0.6 (DOWN) with p-value ≤ 0.05 **(A-C)**, compared to the RNA identified in complex with TIAR immunoprecipitated (RIP) from K562 cells cultured in CO for 72h **(B)** or to the results of rMATS analysis of alternative splicing of cassette exon (CE) and intron retention (RI) **(C)**. Number of genes/proteins in the intersection provided above the bars; group size above the dots.

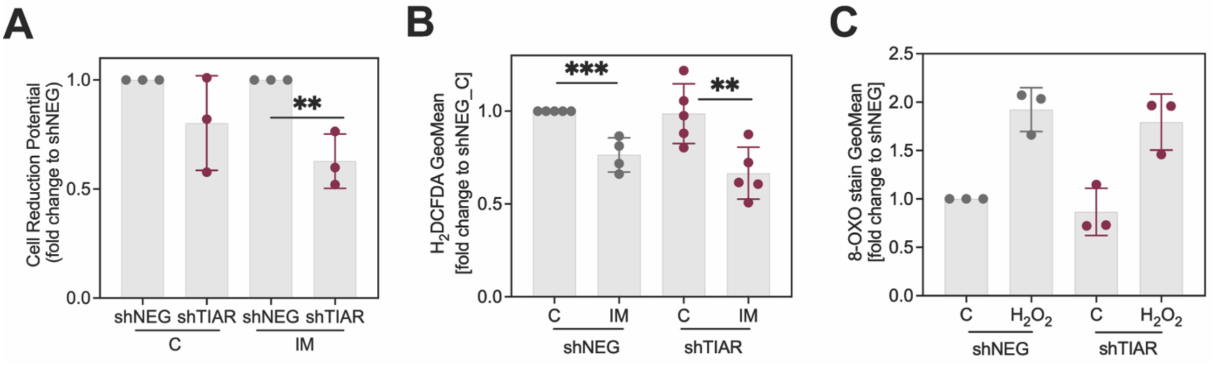

**Figure S14. Impact of imatinib and TIAR knock-down on processes related to oxidation.**

Results from K562 cells with shNEG or shTIAR that were co-cultured with HS-5 cells for 48h before the addition of imatinib (IM) or untreated (C) for 18h. Cell culture and all cell manipulations were at 1.5% $O_2$ in a hypoxia workstation using equilibrated buffers until cell lysis **(A)**, measurement **(B)**, and fixation **(C)**.

**(A)** Cell reduction potential measured using a luminescent MTT assay, normalized to the cell number, expressed as fold change to shNEG (n=3).

**(B)** Level of reactive oxygen species determined in cells using $H_2$DCFDA fluorescent probe and flow cytometry, expressed as fold change to shNEG untreated (n=5).

**(C)** Detection of oxidation-induced DNA/RNA damage using an antibody recognizing 8-hydroxy-2′-deoxyguanosine (8-OXO) conjugated to FITC, analyzed by flow cytometry (n=3) in C and cells treated with 100 µM $H_2O_2$ for 1h.

**(A-C)** Mean value from independent experiments ± SD. Student's two-way t-test was used to compare IM vs C and shTIAR vs shNEG; ** $p \leq 0.01$, *** $p \leq 0.005$.

**Table S1.**

| Figure 6B | | | | |
|---|---|---|---|---|
| Protein DOWN | | | Protein UP | |
| ACADVL | GREM1 | RPL31 | ACOT7 | MINOS1-NBL1 |
| ACTN1 | HEBP1 | S100A4 | ADAM10 | MKI67 |
| ADD2 | HS2ST1 | SARS | AMMECR1 | MTFR1L |
| AGFG1 | HSPA5 | SERPINB2 | ANXA4 | NAA20 |
| AKR1A1 | HSPB1 | SERPINE1 | ATP5H | OPTN |
| ALDH1B1 | IL1A | SLC16A1 | BANF1 | PDAP1 |
| ANKRD10 | IL1B | SLC2A3 | BCAP29 | PIR |
| ANXA1 | IMPA1 | SLC30A1 | BCLAF1 | PITX1 |
| ANXA2 | JUN | SLC38A2 | CASP3 | PPIH |
| ANXA5 | KRT18 | SLC3A2 | CCDC12 | PPM1G |
| ANXA6 | KTN1 | SLC7A5 | CT45A2 | RPS23 |
| APOE | LAPTM5 | SQSTM1 | DCUN1D5 | RRAS |
| ARG2 | LGALS1 | STAU2 | DEPDC1B | RRP15 |
| ARHGEF7 | MMP1 | STX3 | EBP | SARNP |
| ASNS | MTHFD2 | SURF4 | EIF4A2 | SERPINI1 |
| ASPH | MYDGF | TFPI2 | EIF4G3 | SLC25A37 |
| BAIAP2 | NAMPT | TIAL1 | EML2 | SLC35E1 |
| BRD2 | NDRG1 | TPM4 | EXOSC4 | SLC38A5 |
| CALCOCO2 | NDRG2 | TRAM1 | FAM192A | SLC4A1 |
| CAPG | NEU1 | TUBB4A | FASN | SNRPD1 |
| CDC42EP4 | NF1 | TUBB6 | GLB1 | SRP14 |
| CRABP2 | NQO1 | TXNDC5 | HBA1 | TIMM13 |
| CROCC | OSBPL8 | TXNIP | HBE1 | TPM1 |
| CTSB | PACSIN2 | TXNRD1 | HBG1 | TPX2 |
| EFHD2 | PFKM | UMPS | HBG2 | TSTA3 |
| EHD2 | PFKP | UQCRQ | HBZ | TUBB1 |
| EIF1 | PHGDH | VAT1 | HEMGN | USP17L15 |
| ENO2 | PKM | VIM | HIST1H1C | WDR12 |
| ESYT2 | PSAT1 | | HMBS | ZMYND8 |
| EZR | PTRF | | HMGN3 | ZNF512 |
| FAM129B | RAB13 | | HSPA14 | ZNF740 |
| FLNB | RALA | | IPO9 | ZW10 |
| FLNC | RFK | | LCP1 | |
| GFPT1 | RHOC | | LSR | |

| Figure 6D | |
|---|---|
| Common protein DOWN & RIP | Common protein UP & RIP |
| AGFG1 | ACOT7 |
| ANXA1 | AMMECR1 |
| ANXA2 | ANXA4 |
| ANXA5 | BANF1 |
| ARG2 | BCLAF1 |
| CALCOCO2 | CASP3 |
| CRABP2 | CCDC12 |
| EFHD2 | DCUN1D5 |
| EIF1 | DEPDC1B |
| ENO2 | EBP |
| EZR | EIF4A2 |
| GFPT1 | GLB1 |
| HSPB1 | HBA1 |
| IMPA1 | HEMGN |
| LAPTM5 | HMGN3 |
| MTHFD2 | HSPA14 |
| NAMPT | NAA20 |
| NDRG2 | PDAP1 |
| PSAT1 | PIR |
| RALA | PPIH |
| RFK | RPS23 |
| RHOC | RRP15 |
| SLC2A3 | SNRPD1 |
| STAU2 | SRP14 |
| STX3 | TPM1 |
| SURF4 | TUBB1 |
| TFPI2 | ZNF512 |
| TPM4 | |
| TRAM1 | |
| TUBB4A | |
| TXNIP | |
| VAT1 | |

| Figure 6F | | | | |
|---|---|---|---|---|
| Common RIP & CE | | | | Common RIP & RI |
| ABI1 | DMAC2 | KLHL24 | SELENOI | C2orf49 |
| ACADM | DNAJC19 | KMT5A | SKP2 | CCNDBP1 |
| ACP1 | ECHDC1 | LAMTOR3 | SMARCA2 | CLK4 |
| ALG3 | EFR3A | LETMD1 | SNHG1 | DHRS1 |
| ANXA2 | EI24 | MAP3K7 | SNRPG | DYNLL1 |
| APOO | EIF3M | MITD1 | SRSF10 | ERAL1 |
| APTX | EIF4E2 | MLX | SRSF6 | EWSR1 |
| ARMC10 | EXOC1 | MRPS18C | SS18 | FAM133B |
| ASH2L | EXOSC1 | MYB | STARD3NL | FBXO9 |
| ATP6AP2 | EXOSC9 | NDRG2 | STARD4 | GAS5 |
| AURKA | FAM76B | NDUFAF5 | STAU2 | HCFC1R1 |
| BCLAF1 | FGFR1OP2 | NEMP1 | SUGT1 | HNRNPLL |
| BNIP2 | FIP1L1 | NIPA2 | TANK | IDH1 |
| C12orf29 | GCLM | NMU | TERF1 | ISCU |
| C12orf57 | GTPBP10 | NSUN2 | TEX261 | MRPS18C |
| C1orf52 | GYPA | NUF2 | TFPI | MTHFD2 |
| CASP8 | HAUS2 | OCIAD1 | THUMPD2 | PPIA |
| CENPA | HAUS4 | OPA1 | TMEM14B | RPAIN |
| CHEK1 | HMGA1 | OXSR1 | TMEM69 | SEC14L1 |
| CLK4 | HMGCS1 | PARPBP | TRAPPC13 | SRSF5 |
| CNOT2 | HMGN3 | PDCD10 | TTC8 | TCF4 |
| CNOT4 | HMGN4 | PEX2 | UBAP2 | TMEM41B |
| COPZ1 | HNRNPDL | PICALM | UBAP2L | |
| COX20 | HP1BP3 | PPP3CB | UBE2D3 | |
| CPSF6 | HPF1 | PRPF40A | UBQLN1 | |
| CREM | HSF2 | PTPRC | UPF3B | |
| CSNK2A1 | IDI1 | R3HDM1 | USP8 | |
| CTBS | ING3 | RFESD | VHL | |
| DAP3 | INIP | RHOT1 | WSB1 | |
| DCUN1D4 | ISCU | RMDN1 | YIPF1 | |
| DECR1 | ITGB3BP | RNMT | YPEL5 | |
| DEPDC1 | ITM2A | RPL7L1 | ZNF207 | |
| DEPDC1B | KLHDC10 | SAAL1 | ZNF326 | |
| | | | ZRANB2 | |

**Table S1. Protein and transcript changes in CML cells treated with imatinib (IM) in hypoxic co-culture, corresponding to comparisons presented in Figure 6**.

Figure 6B - list of proteins UP or DOWN regulated based on quantitative (QuaNCAT) analysis of protein synthesis in shTIAR versus shNEG. Figure 6D - list of proteins UP or DOWN from QuanCAT for which transcripts were detected in UV-crosslinked RNA immunoprecipitated with TIAR complexes (RIP) from cells in hypoxic co-culture. Figure 6F - list of transcripts detected in RIP, which also had changes in cassette exon inclusion (CE) or intron retention (RI) in shTIAR versus shNEG alternative splicing analysis of RNA from IM-treated cells in hypoxic co-culture.

**Table S2.**

| Gene ID | CCDS ID | Exon No. | F or R | Primer sequence 5'->3' | Amplicon size bp |
|---|---|---|---|---|---|
| *EXOSC5* | CCDS12580.1 | 1 | F | AGGAGGAGACGCATACTGAC | 237 |
| | | 2 | R | TCAGGATCACTTCGAGTGTGG | |
| *HNRNPDL* | CCDS75153.1 | 1 | F | TCCATACAACGCTCCGCCG | 201 |
| | | 2 | R | GCTTGTATCCCAGCTCAAGCC | |
| *SLC25A37* | CCDS47828.1 | 1 | F | GATGGGGACAGCCGAGATG | 145 |
| | | 1 | R | ACCGGGTACATGACCGAGT | |
| *HBE1* | CCDS7756.1 | 1 | F | ATGGTGCATTTTACTGCTGAGG | 155 |
| | | 2 | R | GGGAGACGACAGGTTTCCAAA | |
| *HMBS* | CCDS58186.1 | 3 | F | AGCTTGCTCGCATACAGACG | 156 |
| | | 5 | R | AGCTCCTTGGTAAACAGGCTT | |
| *HSPB1* | CCDS5583.1 | 3 | F | TGGACCCCACCCAAGTTTC | 167 |
| | | 3 | R | CGGCAGTCTCATCGGATTTT | |
| *JUN* | CCDS610.1 | 1 | F | TCCAAGTGCCGAAAAAGGAAG | 129 |
| | | 1 | R | GTTTAAGCTGTGCCACCTGTT | |
| *CFLAR* | CCDS77505.1 | 1 | F | GAAAGAGGTAAGCTGTCTGTCG | 140 |
| | | 1 | R | GTCCGAAACAAGGTGAGGGTT | |
| *AHNAK* | CCDS31584 | 5 | F | GGTAGGCCAGTAGAGGTACAG | 161 |
| | | 5 | R | CCCCACAGAGACTTCAGGT | |
| *SOD1* | CCDS33536.1 | 3 | F | GGTGGGCCAAAGGATGAAGAG | 226 |
| | | 5 | R | CCACAAGCCAAACGACTTCC | |
| *TXNIP* | CCDS72876.1 | 1 | F | TGTGTGAAGTTACTCGTGTCAAA | 107 |
| | | 1 | R | GCAGGTACTCCGAAGTCTGT | |
| *SOD2* | CCDS83141.1 | 3 | F | GGAAGCCATCAAACGTGACTT | 115 |
| | | 3 | R | CCCGTTCCTTATTGAAACCAAGC | |

**Table S2. Sequence of forward (F) and reverse (R) primers used in real-time PCR to quantify cDNA obtained from treverse transcription of RNA from whole cells and immunoprecipitation of TIAR complexes.**

**Table S3.**

| | Read status | | | |
|---|---|---|---|---|
| Sample | Sequenced | Mapped | Unassigned_ NoFeatures | Unassigned_ Ambiguity |
| Sequencing of mRNA from xenografts | | | | |
| mono_R1 | 26814860 | 21697596 | 3200320 | 1916944 |
| mono_R2 | 27162278 | 21236691 | 4104258 | 1821329 |
| mono_R3 | 22070780 | 18531073 | 1805113 | 1734594 |
| **mono_R1-3 sum** | **76047918** | **61465360** | **9109691** | **5472867** |
| shNEG_C_R1 | 17861048 | 15180409 | 1219422 | 1461217 |
| shNEG_C_R2 | 20456664 | 17324147 | 1377236 | 1755281 |
| shNEG_C_R3 | 17827632 | 15029844 | 1193561 | 1604227 |
| **shNEG_C_R1-3 sum** | **56145344** | **47534400** | **3790219** | **4820725** |
| shNEG_IM_R1 | 25380147 | 20484421 | 3187552 | 1708174 |
| shNEG_IM_R2 | 17080368 | 13996674 | 1890704 | 1192990 |
| shNEG_IM_R3 | 25215698 | 21195657 | 2157365 | 1862676 |
| **shNEG_IM_R1-3 sum** | **67676213** | **55676752** | **7235621** | **4763840** |
| shTIAR_C_R1 | 68413058 | 51959527 | 12163085 | 4290446 |
| shTIAR_C_R2 | 16480334 | 14022958 | 1058392 | 1398984 |
| shTIAR_C_R3 | 26843578 | 22817072 | 1848012 | 2178494 |
| **shTIAR_C_R1-3 sum** | **111736970** | **88799557** | **15069489** | **7867924** |
| shTIAR_IM_R1 | 15493295 | 12946481 | 1286968 | 1259846 |
| shTIAR_IM_R2 | 18585073 | 15801693 | 1130257 | 1653123 |
| shTIAR_IM_R3 | 16311011 | 13689956 | 1252047 | 1369008 |
| **shTIAR_IM_R1-3 sum** | **50389379** | **42438130** | **3669272** | **4281977** |

| | Read status | | | |
|---|---|---|---|---|
| Sample | Sequenced | Mapped | Unassigned_ NoFeatures | Unassigned _Ambiguity |
| Sequencing of mRNA from hypoxic co-culture | | | | |
| shNEG_C_R1 | 29251424 | 24107217 | 3268672 | 1875535 |
| shNEG_C_R2 | 18705999 | 15821182 | 1614847 | 1269970 |
| shNEG_C_R3 | 29695204 | 25038139 | 2699992 | 1957073 |
| **shNEG_C_R1-3 sum** | **77652627** | **64966538** | **7583511** | **5102578** |
| shNEG_IM_R1 | 29146359 | 23556565 | 3854879 | 1734915 |
| shNEG_IM_R2 | 29902268 | 24244959 | 3808665 | 1848644 |
| shNEG_IM_R3 | 29450692 | 24460640 | 3062314 | 1927738 |
| **shNEG_IM_R1-3 sum** | **88499319** | **72262164** | **10725858** | **5511297** |
| shTIAR_C_R1 | 32947574 | 27748634 | 3015474 | 2183466 |
| shTIAR_C_R2 | 38712740 | 32365799 | 3917606 | 2429335 |
| shTIAR_C_R3 | 33551222 | 26712479 | 4743035 | 2095708 |
| **shTIAR_C_R1-3 sum** | **105211536** | **86826912** | **11676115** | **6708509** |
| shTIAR_IM_R1 | 26552105 | 22459099 | 2294011 | 1798995 |
| shTIAR_IM_R2 | 47217865 | 36906861 | 7527001 | 2784003 |
| shTIAR_IM_R3 | 43299680 | 35804082 | 4569789 | 2925809 |
| **shTIAR_IM_R1-3 sum** | **117069650** | **95170042** | **14390801** | **7508807** |

**Table S3. Short-read RNA sequencing and mapping statistics.**

Sequenced - number of raw reads sequenced on the Illumina platform; Mapped - counts of mapped reads; Unassigned_NoFeatures - number after filtering out the reads mapped to a region that is not annotated in hg38 annotation; Unassigned_ambiguity - number after filtering for reads mapped to multiple locations in hg38 reference genome; R - biological replicate; C- control cells; IM – imatinib treated.
